# Imbalanced immune response and dysregulation of neural functions underline fatal opportunistic encephalitis caused by astrovirus

**DOI:** 10.1101/2022.08.20.504643

**Authors:** Olga A. Maximova, Melodie L. Weller, Tammy Krogmann, Daniel E. Sturdevant, Stacy Ricklefs, Kimmo Virtaneva, Craig Martens, Kurt Wollenberg, Mahnaz Minai, Ian N. Moore, Craig S. Sauter, Juliet N. Barker, W. Ian Lipkin, Danielle Seilhean, Avindra Nath, Jeffrey I. Cohen

## Abstract

The incidence of infections of the central nervous system (CNS) in humans is increasing due to emergence and reemergence of pathogens and an increase in the number of immunocompromised patients. Many viruses are opportunists and can invade the CNS if the immune response of the host is impaired. Here we investigate neuropathogenesis of a rare CNS infection in immunocompromised patients caused by astrovirus and show that it shares many features with another opportunistic infection of the CNS caused by human immunodeficiency virus. We show that astrovirus infects CNS neurons with a major impact on the brainstem. In the setting of impaired peripheral adaptive immunity, host responses in the astrovirus infected brain are skewed to the innate immune response with exuberant activation of microglia and macrophages. Astrovirus infection of neurons and responses by phagocytic cells lead to disrupted synaptic integrity, loss of afferent innervation related to infected neurons, and global impairment of both excitatory and inhibitory neurotransmission. The response employed in the CNS against opportunistic viruses, such as astrovirus and HIV, may be a common compensatory defense mechanism which inadvertently leads to loss of neural functions due to the host’s exuberant innate immune response to pathogens when adaptive immunity is impaired.

## INTRODUCTION

Human CNS infections are increasing due to spread of pathogens to new areas, emergence of new pathogens or an increase in vulnerable populations [1] such as persons receiving immunosuppressive medications, immune senescence associated with longer lifespans, or infection with viruses inducing an immunodeficient state [2]. Many viruses, including West Nile virus, Eastern equine virus, human herpesvirus 6, enteroviruses, and JC virus invade the CNS when infection is poorly controlled due to impaired immunity of the host [3]. One group of viruses that has recently been reported to cause opportunistic CNS infections is the astroviruses (AstVs). AstVs are RNA viruses that infect the gastrointestinal tract of mammals and birds and cause fever, nausea, and diarrhea in humans [4]. Three clades of astroviruses infect humans; HAstV1 to HAstV8 are the classical human strains, while HAst-MLB1-3 and HAst-VA1-5 have been reported more recently [5]. These newer clades are more genetically related to other animal astroviruses than they are to classical human strains. Gastroenteritis due to astroviruses is more common in children, especially those under 2 years old. Outbreaks have occurred in the elderly and in immunocompromised patients. The latter can have chronic diarrhea with prolonged shedding of virus. Ten cases of encephalitis due to astroviruses have been reported in humans [6-14]. In most cases, astrovirus encephalitis developed in the immunocompromised patients [6-13] and was fatal [6, 7, 9, 10, 12, 13]. Only two cases were reported in apparently immunocompetent patients who survived the disease [13, 14]. Astrovirus has also been associated with neurologic disease in animals [15].

The pathogenesis of CNS disease due to neuroinvasive astrovirus infection in humans is not well understood. Cellular targets of astroviruses in the human CNS remain elusive since astrovirus capsid protein or RNA have been reported only in three studies as being present in both neuronal and glial cells [6, 9, 10]. Host responses to astrovirus infection in the human CNS and impact of this infection on neurophysiology have not been comprehensively studied.

Here, we investigated transcriptional changes in the CNS of three patients with astrovirus neurological disease (AstV-ND) and compared the results with another opportunistic viral infection of the CNS, human immunodeficiency virus (HIV-1) associated neurocognitive disorder (HAND). Further analysis of protein expression and affected cell types in the brain from a patient with AstV-ND revealed specific cellular targets of the virus, domination of host responses by the microglia and macrophages, and impairment of specific neural functions.

## RESULTS

### Identification of a neuroinvasive human astrovirus (HAstV-NIH) closely related to other neuropathogenic astroviruses detected in humans

We investigated the cause of encephalitis in a 58-year-old-man who had undergone umbilical cord blood transplant for lymphoma and received immunosuppression for graft-versus-host disease (see Materials and Methods for the case report). RNA from the patient’s brain was hybridized to a virus chip containing over 3000 probes for viral families and astrovirus was detected (Supplemental Fig. 1a). To confirm this finding, RNA was isolated from a different portion of the patient’s brain, cDNA was prepared, and PCR was performed using pan astrovirus primers to the polymerase gene of the virus. Astrovirus sequences were detected in the patient’s brain tissue, but not in control human brain (Supplemental Fig. 1b). In situ hybridization demonstrated that brain tissue from the patient, but not from a control brain, hybridized to the astrovirus probe (Supplemental Fig. 1, c and d). The newly detected virus was designated as human astrovirus HAstV-NIH, and the case of astrovirus neurological disease associated with this virus was designated as AstV-ND-1.

Sequencing of the capsid gene from patient AstV-ND-1 followed by phylogenetic analysis placed the virus together with four human astroviruses detected in the brain from previously reported cases of encephalitis in immunocompromised patients (3 pediatric [6, 8, 10] and one adult [9] cases, Supplemental Fig. 1e). These astroviruses were: HAst-V-PS ([6] patient referred to as AstV-ND-2 in this manuscript), HAstV-VA1/HMO-C-PA ([8] patient referred to as AstV-ND-3 in this manuscript), and two viruses from VA/HMO clade (HAstV-VA1/HMO-C-UK1) [9, 10]. Together, these five neuroinvasive astroviruses form a separate group from the astrovirus of the MLB1 lineage (HAst-MLB1 NAGANO 1545) detected in an immunocompromised boy with acute encephalopathy (Supplemental Fig. 1e [11]) and from the other classical human strains (i.e., HAstV 2, 5, and 8). The closest relatives to the human neuroinvasive astroviruses are eight bovine astroviruses linked to encephalitis and one ovine astrovirus (Supplemental Fig. 1e).

**Supplemental Figure 1.**
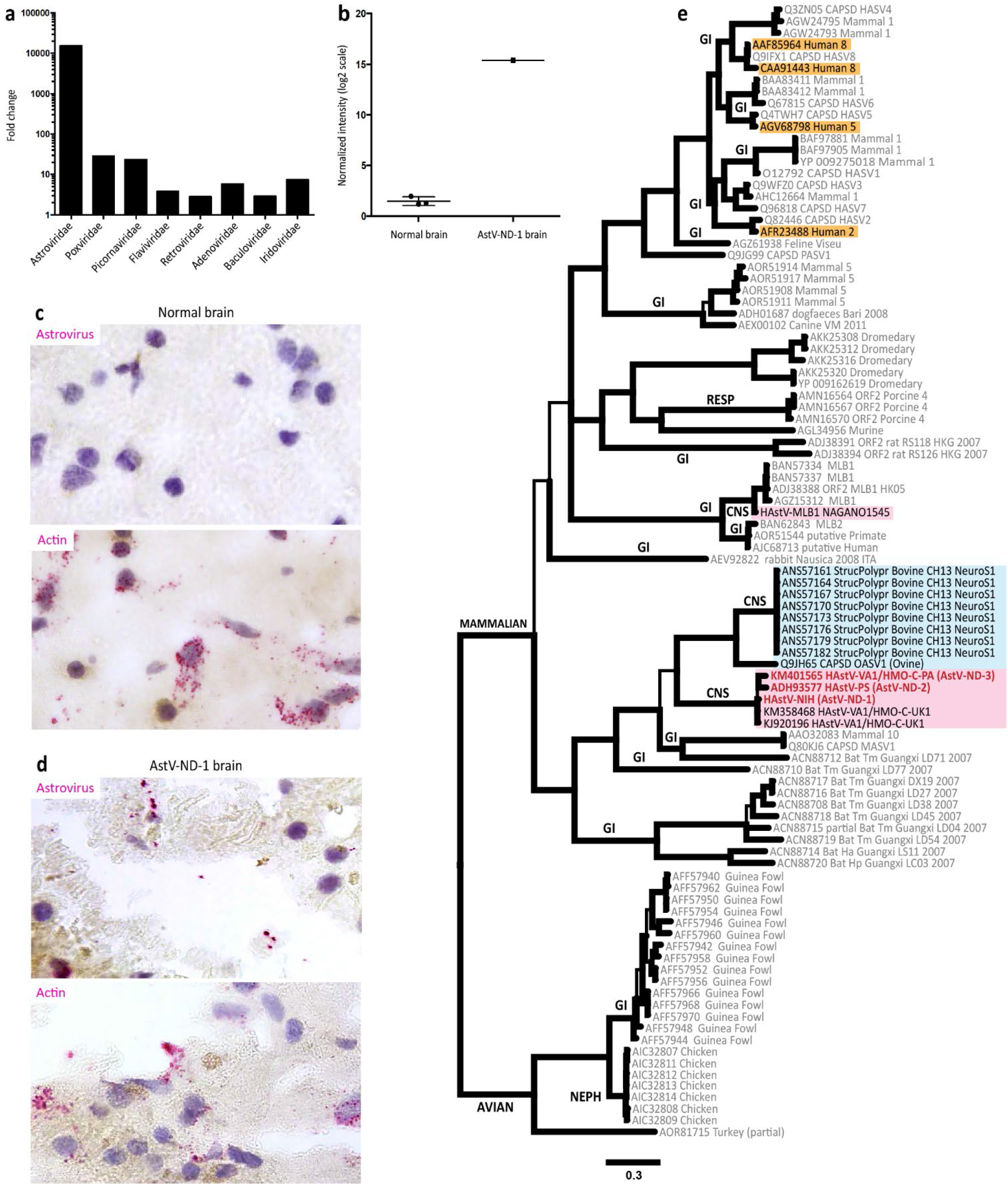
Detection and phylogenetic analysis of a new neuroinvasive astrovirus HAstV-NIH causing encephalitis. (**a**) Fold change relative to control for probes to virus families hybridizing to RNA from the AstV-ND-1 patient’s brain. (**b**) Astrovirus RNA detection by PCR in the brain of patient AstV-ND-1. (**c** and **d**) In situ hybridization signals (magenta-red) in indicated brain tissue samples using the negative strand astrovirus RNA probe (upper panels) or RNA probes for actin (lower panels). (**c**) Bayesian phylogenetic tree of 97 Astroviridae ORF2/capsid protein sequences. Thick branches are significantly supported (posterior probability > 0.95) by the data. Mammalian and avian lineages are labeled on their most-basal branches. Known clinical presentations are indicated (CNS, associated with CNS disease; GI, gastrointestinal; RESP, respiratory; and NEPH, nephritic). Six astroviruses associated with CNS infection are highlighted in magenta. Classical human astroviruses are highlighted in orange. Mammalian astroviruses closely related to five human neuroinvasive astroviruses are highlighted in cyan. Astroviruses in brain samples from three AstV-ND cases (AstV-ND-1, AstV-ND-2, and AstV-ND-3) investigated in this study are indicated in red font.

### CNS host defense response to astrovirus infection in immunocompromised hosts is skewed to the innate immune response

We first used an agnostic approach to gain a broad insight into the pathogenesis of astrovirus infection in the CNS, by analyzing global changes in the brain transcriptome of patients with AstV-ND. RNA-seq was performed on brain samples of three patients with AstV-ND: (i) AstV-ND-1, postmortem sample, unknown brain site (see Materials and Methods for case report), (ii) AstV-ND-2 (a 15-year-old boy with X-linked agammaglobulinemia [XLA]) [6], postmortem sample of the brainstem), and (iii) AstV-ND-3 (a 14-year-old boy with XLA) [8], biopsy sample from the frontal cortex). Since brain samples from these cases were from different (or unknown) anatomical sites, we added for comparison five major anatomical sites (i.e., frontal cortex, thalamus, brainstem, medulla [part of the brainstem], and cerebellum) from subjects without known neurological disease (normal controls).

We performed a functional enrichment analysis and used the Panther statistical enrichment test (SET) [16] on unnormalized gene expression data in each sample separately. SET accepts all expressed genes and their expression values (i.e., number of reads > 0, for this study) to identify significantly enriched gene ontology (GO) terms for genes whose expression has shifted to the larger values (large shifts) or smaller values (subtle shifts) compared to the overall distribution of values. SET determines intrinsic biological processes that are under selective transcriptional control in a given tissue and tissue compartments where these biological processes act (i.e., within the cell and/or extracellular space). This analysis allowed for unbiased probing for coordinated shifts in global gene expression without the need to normalize data by choosing specific pairs of diseased and normal tissues to calculate DEGs.

The SET identified large coordinated transcriptional shifts that confirmed the CNS origin of both normal and AstV-ND tissue samples (Fig. 1, a - d; black background). These large shifts were indicative of neurophysiological processes occurring in the specific cellular compartments of the CNS such as neuronal cell bodies, dendrites, axons, synapses (including glutamatergic [excitatory] and GABA-ergic [inhibitory] synapses), and glial cells and their projections (Fig. 1, a - d; indicated by black text).

**Figure 1.**
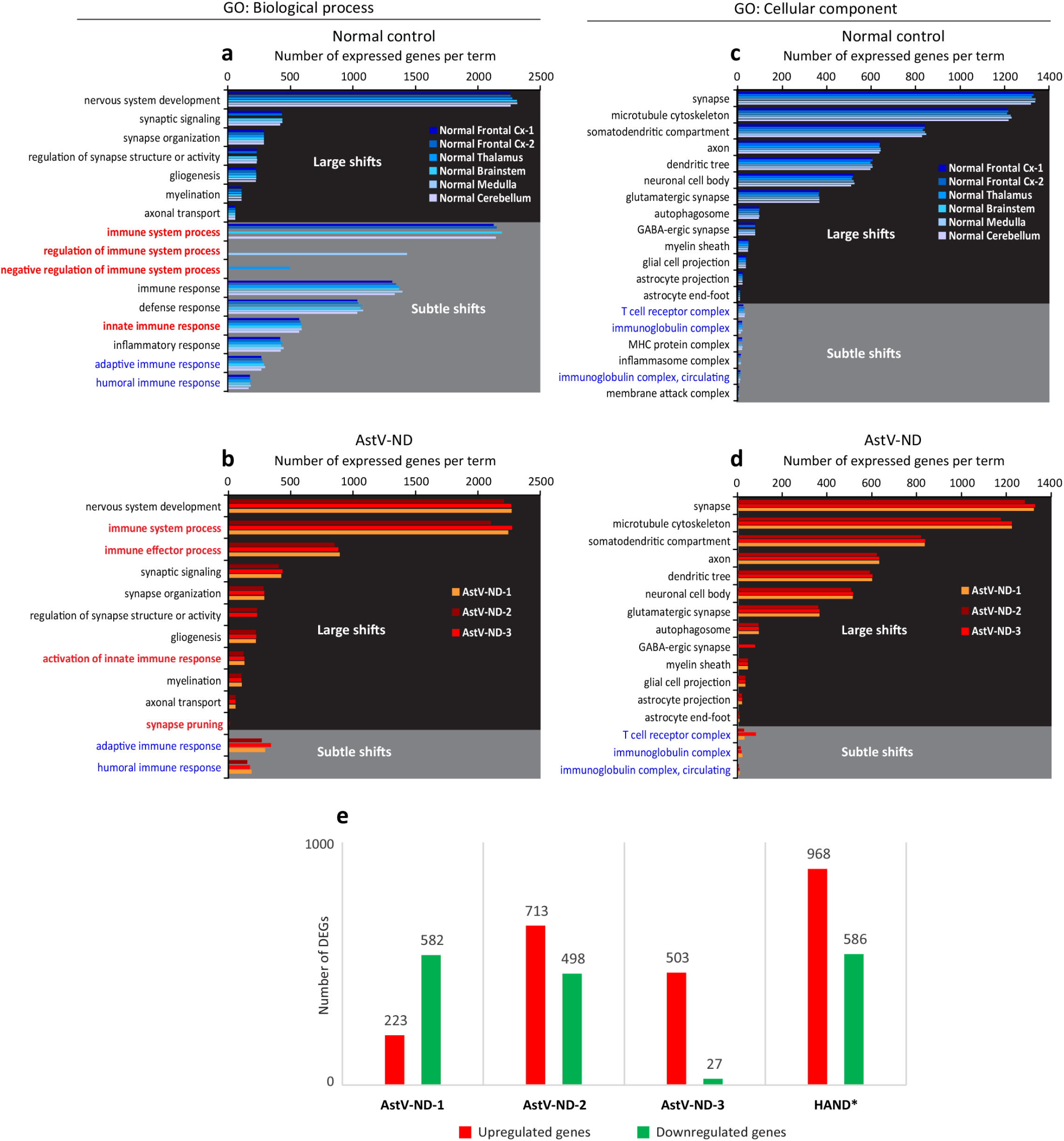
Changes in global gene expression in the brain due to astrovirus infection. (**a** - **d**) Graphs show selected significantly enriched GO terms (FDR < 0.05) and numbers of genes expressed per GO term, as determined by the Panther SET based on unnormalized gene expression data in six normal human brain samples (**a** and **c**) and one brain sample from each of indicated AstV-ND cases (**b** and **d**). Large coordinated transcriptional shifts (i.e., shifts to the greater number of reads from the overall distribution) are indicated by black background and subtle coordinated transcriptional shifts (i.e., shifts to smaller number of reads from the overall distribution) are indicated by gray background. Biological functions of intertest that switched from the subtle shifts in normal brains to large shifts in AstV-ND brains (including emergence of synapse pruning) are highlighted by red text in **a** and **b**. GO terms of interest describing the transcriptional control of the biological functions (**a** and **b**) and cellular/extracellular components (**c** and **d**) that remained subtle in both normal and AstV-ND brains are highlighted by blue text in **a** - **d**. (**e**) Numbers of DEGs in three cases of AstV-ND and previously published average number of DEGs in brain samples (frontal lobe white matter) from five patients (ages: 43-58 years old) with HIV-1-associated neurocognitive disorders (HAND) [17]).

Subtle coordinated transcriptional shifts in the normal brain regions were associated with immune processes (including innate and adaptive immune responses) (Fig. 1a; gray background) and immune system complexes (Fig. 1c; gray background). This is consistent with known tight control of the immune and inflammatory processes in normal human CNS. Complete SET results for normal brain regions are provided in Supplemental File 1.

In the brain of all three patients with AstV-ND, transcriptional control of the immune system processes switched from subtle to large coordinated shifts, with a change of the innate immune processes to an activated state, and initiation of the synaptic pruning (compare normal control and AstV-ND Fig. 1, a and b, respectively; highlighted by red text). However, in contrast to activation of the innate immune responses, transcriptional regulation of the adaptive and humoral immune responses (as well as associated immune system complexes) in the brain of patients with AstV-ND did not exhibit either activation and/or shifts to larger expression values (compare normal control and AstV-ND in Fig. 1, a - c; highlighted by blue text). Complete SET results for AstV-ND brain samples are provided in Supplemental File 2.

These results suggest that the CNS host response to neuroinvasive astrovirus infection in patients with deficiencies in adaptive immunity are skewed to the innate immune response.

### Functional patterns of gene upregulation in the brain of patients with astrovirus or HIV infection are dominated by microglial and macrophagic signatures

We next determined which genes were upregulated in the brain of patients with astrovirus encephalitis and compared the results with those previously reported from the brains of five patients with another opportunistic brain infection - human immunodeficiency virus-associated neurocognitive disorder (HAND). The patients with untreated HAND (HAND 1 - 5; Table 1 in reference [17]) had low CD4 counts and high levels of HIV in the brain.

**Table 1.** Attributes of astrovirus infections of the CNS in humans and animals.

| Underlying disease/<br>Immunosuppressive treatments | Infected CNS<br>regions/CSF<sup>b</sup> | Infected<br>cells | Neuropathology/diagnosis | Outcome | Reference |
| --- | --- | --- | --- | --- | --- |
| **Humans** |  |  |  |  |  |
| Diffuse large B cell lymphoma<br>Immunosuppressive treatments<sup>a</sup> | - Brainstem<br>- Cerebellum<br>- Thalamus<br>- Basal ganglia<br>- Hippocampus<br>- Cerebral<br>cortex | Neurons | - microglial activation and perineuronal topology;<br>- infiltration by peripheral macrophages;<br>- moderate astrogliosis;<br>- limited infiltration by T cells;<br>- absence of infiltration by B cells;<br>- transcriptional dysregulation of neural functions;<br>- disruption of synaptic integrity;<br>- loss of afferent innervation of infected neurons;<br>- impairment of excitatory/inhibitory neurotransmission | fatal | this study |
| X-linked agammaglobulinemia | Brainstem<sup>c</sup><br>Cerebellum<br>Frontal cortex<sup>d</sup><br>Basal ganglia | Astrocytes<br>Neurons | - astrogliosis;<br>- microgliosis;<br>- perivascular/parenchymal infiltration by T cells and<br>macrophages;<br>- neuronal loss;<br>- axonal swelling;<br>- myelin degeneration | fatal | [6]<br>and this<br>study |
| Severe primary combined<br>immunodeficiency | Brainstem<br>Cerebellum<br>Basal ganglia<br>Hippocampus | not reported | - leptomeningeal/periventricular infiltration by macrophages<br>- neuronal degeneration | One of 3<br>cases<br>was fatal | [7] |
| Chronic lymphocytic leukemia<br>Immunosuppressive treatments | Frontal cortex | Neurons | - astrogliosis<br>- microgliosis<br>- minimal parenchymal infiltration by T cells | fatal | [9] |
| X-linked agammaglobulinemia | Frontal cortex | not reported | - microgliosis<br>- perivascular/parenchymal infiltration by T cells and<br>macrophages | recovery | [8]<br>and this<br>study |
| Acute myeloid leukemia<br>Immunosuppressive treatments | CSF | not reported | Meningitis | fatal | [13] |
| Congenital aplastic anemia<br>Immunosuppressive treatments | CSF | not reported | Encephalopathy | recovery | [11] |
| Cartilage hair hypoplasia<br>and associated immunodeficiency<br>Immunosuppressive treatments | Cerebral cortex | Neurons | - degeneration of neurons<br>- robust microgliosis/ macrophage infiltration<br>- astrogliosis<br>- minimal parenchymal infiltration by T cells | fatal | [10] |
| Acute myeloid leukemia<br>Immunosuppressive treatments | Unspecified | not reported | - astrogliosis<br>- perivascular/leptomeningeal inflammatory cell infiltration | fatal | [12] |
| Immunocompetent | CSF | not reported | Meningitis | recovery | [13] |
| Immunocompetent | CSF | not reported | Encephalopathy | recovery | [14] |
| **Cattle** |  |  |  |  |  |
| unknown | Cerebellum<br>Brainstem<br>Spinal cord<br>Midbrain<br>Thalamus<br>Basal ganglia<br>Cerebral cortex | Neurons | - degeneration of neurons<br>- microgliosis<br>- astrogliosis (Bergman glia)<br>- perivascular infiltration (lymphocytes, macrophages, and<br>plasma cells) | Necropsy<br>report | [26] |
| unknown | Brainstem<br>Cerebellum<br>Cerebrum<br>Hippocampus | Neurons | - degeneration of neurons<br>- microgliosis<br>- perivascular infiltration | Necropsy<br>report | [38, 39] |
| unknown | Medulla | Neurons | - degeneration of neurons<br>- microgliosis | Necropsy<br>report | [40] |
| **Pigs** |  |  |  |  |  |
| unknown | Spinal cord<br>Brainstem<br>Cerebellum<br>Cerebral cortex | Neurons | - degeneration of neurons<br>- microgliosis<br>- perivascular/leptomeningeal infiltration | Necropsy<br>report | [41] |

<sup>a</sup> B cell depleting and T cell immunosuppressive therapy (rituximab, cyclophosphamide, doxorubicin, vincristine, prednisone, dexamethasone, cytarabine, and cisplatin for treatment of lymphoma); myeloablative conditioning therapy (cyclophosphamide, thiotepa, fludarabine, and total body irradiation); umbilical cord blood stem cell transplant; graft-versus-host disease prophylaxis and treatment (cyclosporine, mycophenolate mofetil, methylprednisolone, and budesonide); and additional B cell depleting therapy (rituximab).
<sup>b</sup> Based on the data from this study or referenced reports, we list the CNS regions from the most to the least affected.
<sup>c</sup> Brainstem postmortem specimen had the highest viral load, followed by the cerebellum and frontal cortex. Frozen brainstem specimen was used in this study for RNA-sequencing.
<sup>d</sup> Frontal cortex biopsy and postmortem specimens were used to detect astrovirus infected cells.

Of the three AstV-ND cases we studied, AstV-ND-1 had the least number of upregulated genes (n=223), followed by AstV-ND-3 (n=503), and AstV-ND-2 (n=713) (Figure 1e). In contrast, the number of upregulated genes in HAND (n=968) was higher compared to the AstV-ND cases. Since the differences in the numbers of DEGs do not necessarily reflect functional differences controlled by the groups of genes acting in certain biological pathways and processes, we performed comparative functional enrichment analyses of upregulated genes in the brain of patients with AstV-ND and HAND. The functional genomics platform gProfiler [18] was used to determine and compare functional enrichments for DEGs based on the simultaneous interrogation of multiple gene ontology domains: molecular function (MF), biological process (BP), cellular component (CC), Kyoto Encyclopedia of Genes and Genomes (KEGG), Reactome (REAC), WikiPathways (WP), TRANSFAC transcriptional factors regulatory motif matches (TF), miRNA targets from miRTar-Base (MIRNA), tissue specificity based on Human Protein Atlas (HPA), protein complexes (CORUM), and Human Phenotype Ontology (HP).

AstV-ND-3 had the most significant upregulation of the immune responses (including innate and adaptive immune responses) to virus infection in the brain compared with the other two AstV-ND cases (Fig. 2c vs. Fig. 2a and 2b; Fig. 3: terms #1, #2, and #6; complete results of functional enrichment analysis are provided in Supplemental File 3). Of note, AstV-ND-3 was the only one of the three patients with astrovirus encephalitis that survived [8]. AstV-ND-2 and AstV-ND-3 were similar in their repertoire of upregulation of immune responses (Fig. 2b and 2c), although the adjusted p values for enrichment of innate and adaptive immune responses were lower in AstV-ND-2 (Fig. 3). As expected due to their underlying primary immunodeficiency (XLA), no significant upregulation of the processes related to immunoglobulin mediated/humoral immune response and B cell mediated immunity was detected in these cases (Fig. 3; terms #26 - 28) and only these two patients had a significant enrichment of the KEGG pathway term “Primary immunodeficiency” (Fig. 2 and Fig. 3; term #37). These findings are consistent with the genetic deficiency in B cell immunity due to XLA and suggest impaired adaptive immune response to astrovirus infection in the brain.

**Figure 2.**
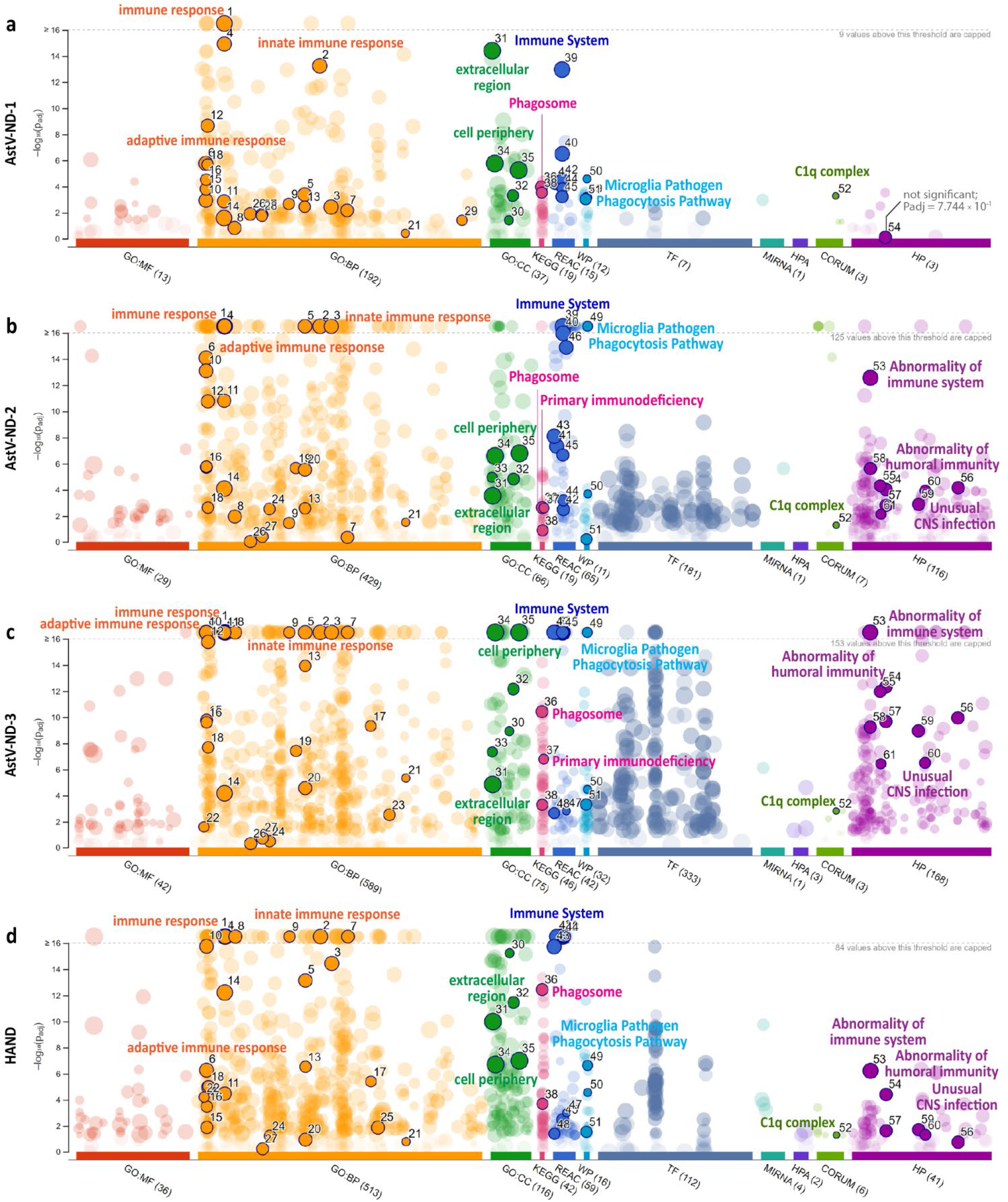
Multi-source visualization of functional enrichment for upregulated gene expression in AstV-ND and HAND. (**a** - **d**) Manhattan plots show significantly enriched terms (P value adjusted [Padj] < 0.05) across multiple ontologies for genes that were upregulated in cases of astrovirus and HIV CNS infections. Circles represent functional terms and are color-coded according to the source ontologies. X-axes show source ontologies. Y-axes show Padj enrichment values (corrected for multiple testing) in a negative log_10_ scale. Padj values above the threshold indicated by gray dashed line are capped, and the number of capped values is indicated. Circle sizes are in accordance with the corresponding term size. The terms of interest are shown by solid colors, outlined (*black border*), and enumerated (the rest of enriched terms appear as semi-transparent circles). The number next to the source name in the x-axis labels shows the number of significantly enriched terms identified from a given source. Select top terms of interest are indicated in each plot by the text colored according to the source ontologies. *Gene ontology data source abbreviations:* MF, molecular function; BP, biological process; CC, cellular component; KEGG, Kyoto Encyclopedia of Genes and Genomes; REAC, Reactome; WP, WikiPathways; TF, TRANSFAC transcriptional factors regulatory motif matches; MIRNA, miRNA targets from miRTar-Base; HPA, tissue specificity based on Human Protein Atlas; CORUM, protein complexes; and HP, Human Phenotype Ontology.

**Figure 3.**
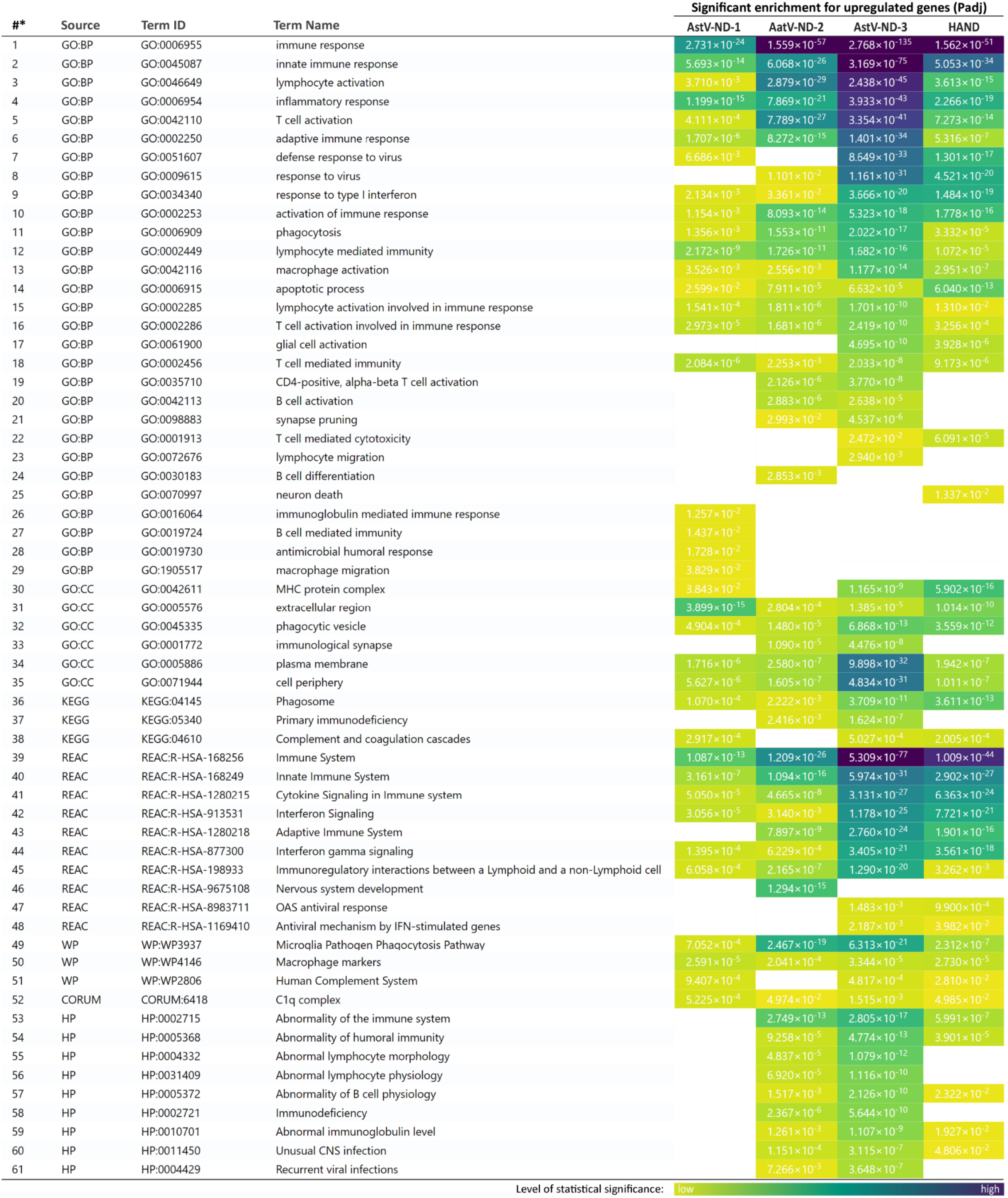
Multi-source side-by-side heatmap of the statistical significance for functional enrichment related to upregulated gene expression in AstV-ND and HAND. Multi-source functional enrichment terms of interest and their corresponding Padj enrichment values are shown for cases of astrovirus and HIV CNS infections. Blanks indicate the absence of significance (Padj > 0.05).* Indicates the term enumeration corresponding to that shown in Fig. 2 (solid color circles). Enrichment values are colored according to the level of statistical significance (scale is shown at the bottom of the figure).

In contrast, AstV-ND-1 patient (with diffuse large B cell lymphoma who had been treated with rituximab, an umbilical cord blood transplant, and corticosteroids for graft-versus-host disease; see case report in the Materials and Methods) had significant, although low level, transcriptional upregulation of B cell mediated immunity and humoral/immunoglobulin mediated immune responses (Fig. 3; terms #26 - 28). However, unlike the two patients with a long-term (about 15 year) immunodeficiency due to XLA, none of the Human Phenotype (HP) ontology terms were significantly enriched in AstV-ND-1, most likely reflecting a relatively shorter-term immunosuppression due to therapies for hematological malignancy. The lower levels of statistical significance of enriched terms for upregulated genes in AstV-ND-1 compared to AstV-ND-2 and AstV-ND-3 (Fig. 3) most likely reflects the lower total number of upregulated genes but may also be related to different responses due to the older age of AstV-ND-1 patient (58 years old) versus AstV-ND-2 and AstV-ND-3 patients (14 and 15 years old).

Functional patterns of gene upregulation in the brain of patients with HAND were more similar to AstV-ND cases with XLA (AstV-ND-2 and AstV-ND-3) than to AstV-ND-1. Of note, all these patients lacked significant enrichment for terms representing B cell immunity and humoral/immunoglobulin mediated immune responses (Fig. 3; terms #26 - 28). Interestingly, five HP terms (Fig. 3; term #53 “Abnormality of the immune system”; term #54 “Abnormality of humoral immunity”; term #57 “Abnormality of B cell physiology”; term #59 “Abnormal immunoglobulin level”; and term #60 “Unusual CNS infection”) were significantly enriched in HAND, similar to AstV-ND-2 and AstV-ND-3, but different from AstV-ND-1. These findings suggest that the long-term immunodeficiency underlying HAND as well as AstV-ND-2 and AstV-ND-3 cases predisposes to a similar transcriptional dysregulation and phenotypic abnormalities in contrast to a shorter-term iatrogenic immunodeficiency.

Functional terms related to the responses by macrophages/microglia (Fig. 3; term #11 “phagocytosis”; term #13 “macrophage activation”; term #36 “Phagosome”; term #49 “Microglia Pathogen Phagocytosis Pathway”; term #50 “Macrophage markers”; and term #52 “C1q (complement component) complex” were significantly enriched in all AstV-ND and HAND cases. These findings and the observation that the “innate immune response” (Fig. 2; Fig. 3, term #2) was the top upregulated biological process related to the immune system process, suggest that transcriptional upregulation of the CNS innate immune responses to these opportunistic infections is robust, but that regulation of adaptive/humoral responses is impaired.

Comparison of the groups of genes that were upregulated in AstV-ND and HAND showed only 32 genes in common (Fig. 4a, all genes associated with the Venn diagram are listed in Supplemental File 4). Analysis of these common upregulated genes revealed a concise functional signature that showed a shared repertoire of the immune responses between these opportunistic neuroinvasive virus infections. The innate immune response with functions related to microglia and macrophages (e.g., term # 11 “Microglia Pathogen Phagocytosis Pathway”; term # 10 “Phagosome”; term #12 “Complement activation”; and term #5 “Synaptic pruning”) ranked consistently higher than the adaptive immune response (Fig. 4, b and c; complete list of significantly enriched terms corresponding to Fig. 4b can be found in Supplemental File 5).

**Figure 4.**
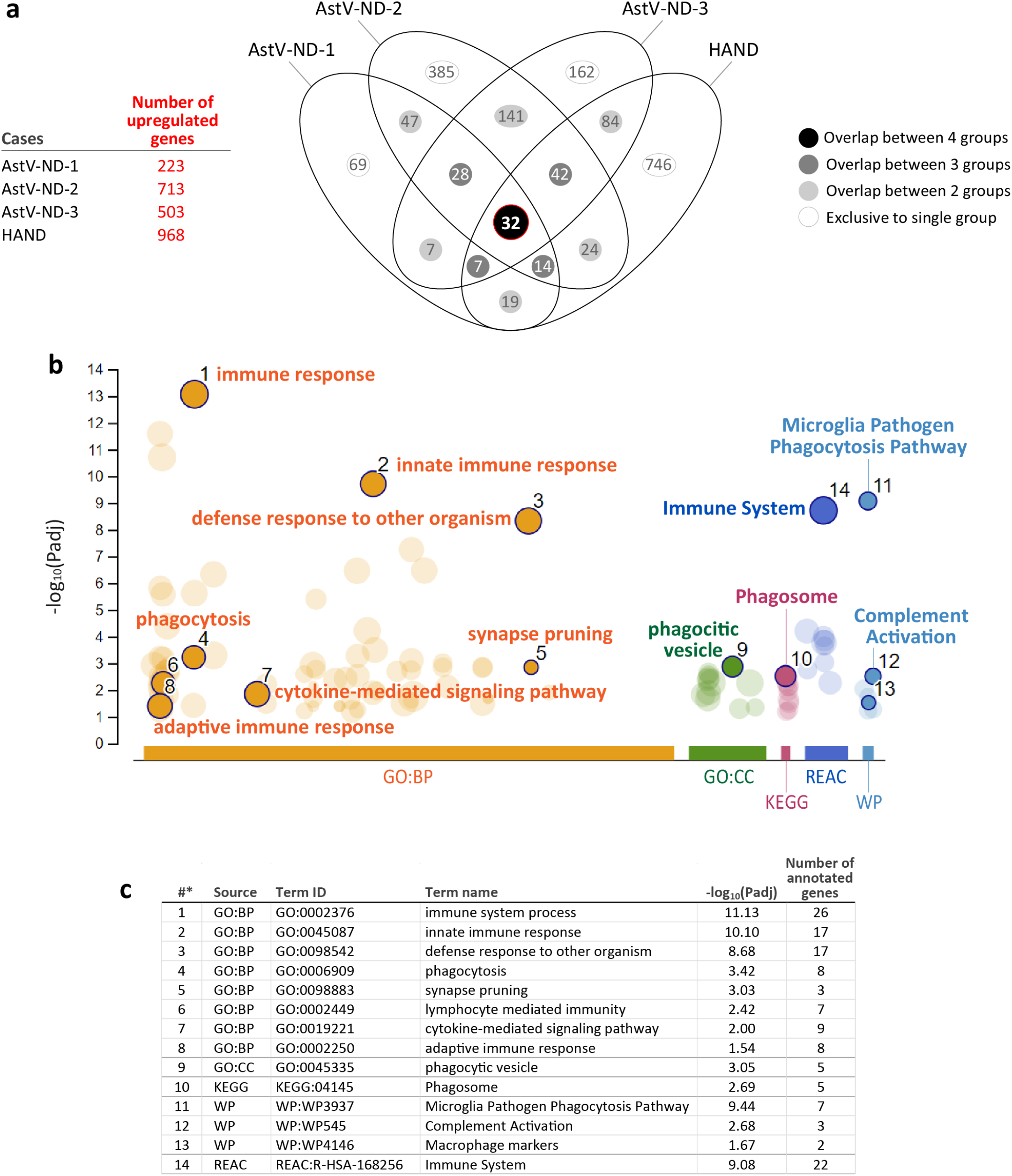
Functional genomic analysis of common upregulated genes in AstV-ND and HAND. (**a**) Venn diagram comparison of the groups of genes that were upregulated in indicated cases of astrovirus and HIV CNS infections. (**b**) Manhattan plot shows significantly enriched terms across indicated ontologies (x-axis) for genes that were commonly upregulated in all cases of AstV-ND and HAND (overlap between 4 groups is highlighted by red border in **a**; n = 32). Circles represent significantly enriched functional terms and are color-coded according to the source ontologies. X-axis: source ontologies. Y-axis: Padj enrichment values in negative log_10_ scale. Circle sizes are in accordance with the corresponding term size. The terms of interest are shown by solid colors, outlined (*black border*), and enumerated (the rest of enriched terms appear as semi-transparent circles). Selected top terms of interest are indicated in each plot by the text colored according to the source ontologies. (**c**) Statistical details for the enriched terms enumerated (denoted as #*) and shown by solid circles in **b**. Gene ontology data source abbreviations are listed in Fig. 2.

Together, these findings show that shared functional patterns of gene upregulation in the brain in immunocompromised persons in response to opportunistic virus CNS infections (AstV-ND and HAND) are dominated by innate immune responses with microglial and macrophagic signatures whereas regulation of the adaptive/humoral responses is impaired.

### Astrovirus and HIV infection of the CNS share functional patterns of gene downregulation that signify a negative impact on neural functions

Having identified shared and unique functional patterns of gene upregulation in the brain of patients with AstV-ND or HAND, we next analyzed genes that were downregulated in the brain of these patients. Among three AstV-ND patients, AstV-ND-1 had the largest number of downregulated genes (n=582), followed by AstV-ND-2 (n=498), and AstV-ND-3 (n=27) (Fig. 1e). The number of downregulated genes in HIVE (n=586) was similar to AstV-ND-1. Unlike AstV-ND-3, AstV-ND-2 had significantly enriched terms based on downregulated genes with functions in the cellular compartments of neurons: “synapse” (term #14); “axon” (term #15); “somatodendritic compartment” (term #16); and “GABA receptor Signaling” (term #31) (Fig. 5 and Fig. 6; complete results are provided in Supplemental File 6).

**Figure 5.**
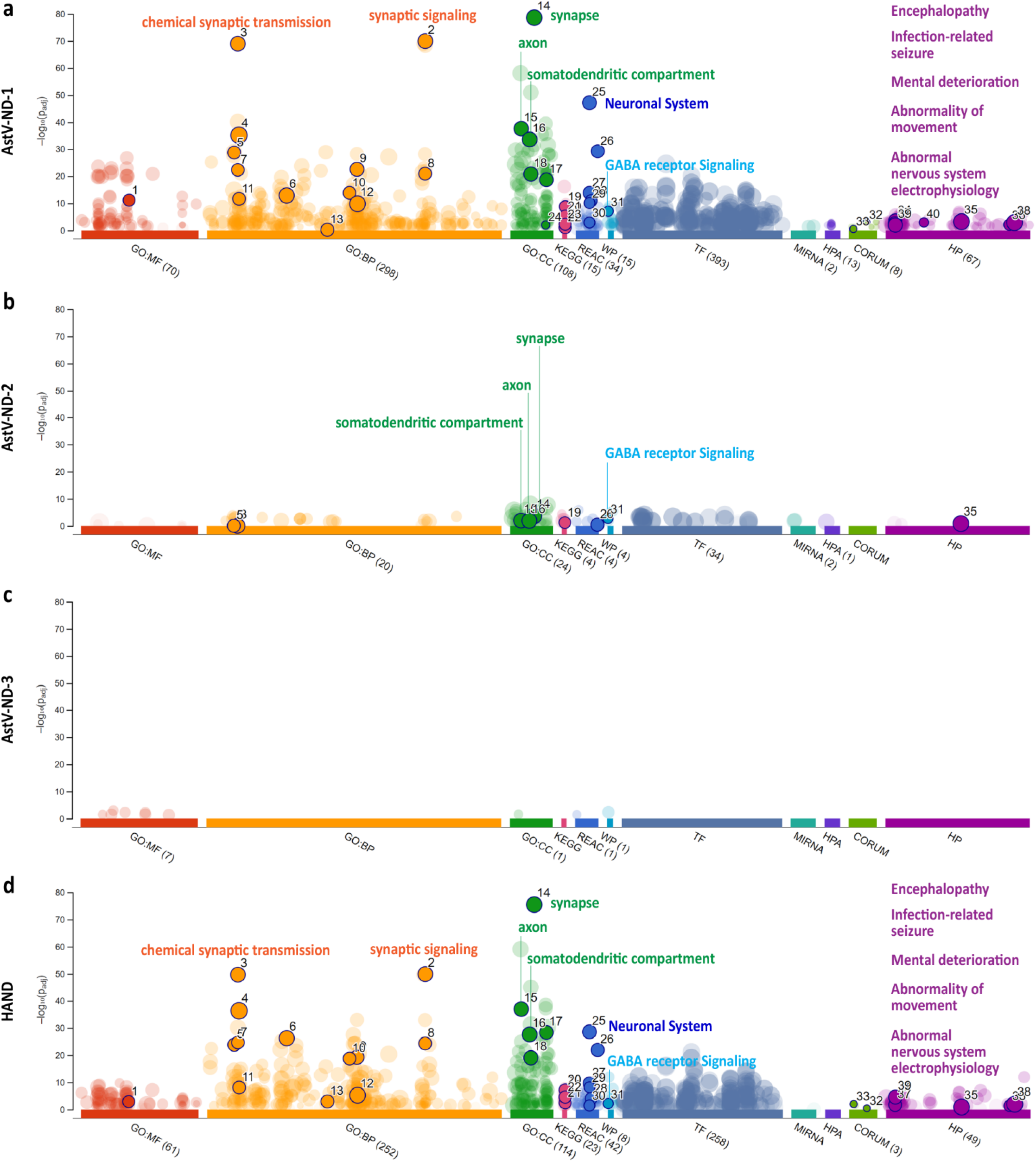
Multi-source visualization of functional enrichment for the downregulated gene expression in AstV-ND and HAND. (**a** - **d**) Manhattan plots simultaneously show significantly enriched terms (Padj < 0.05) across multiple ontologies for genes that were downregulated in cases of astrovirus and HIV CNS infections. Circles represent functional terms and are color-coded according to the source ontologies. X-axes show source ontologies. Y-axes show Padj enrichment values (corrected for multiple testing) in a negative log_10_ scale. Padj values above the threshold indicated by gray dashed line are capped and the number of capped values is indicated. Circle sizes are in accordance with the corresponding term size. The terms of interest are shown by solid colors, outlined (*black border*), and enumerated (the rest of enriched terms appear as semi-transparent circles). The number next to the source name in the x-axis labels shows the number of significantly enriched terms identified from a given source. Select top terms of interest are indicated in each plot by the text colored according to the source ontologies. Gene ontology data source abbreviations are listed in Fig. 2.

**Figure 6.**
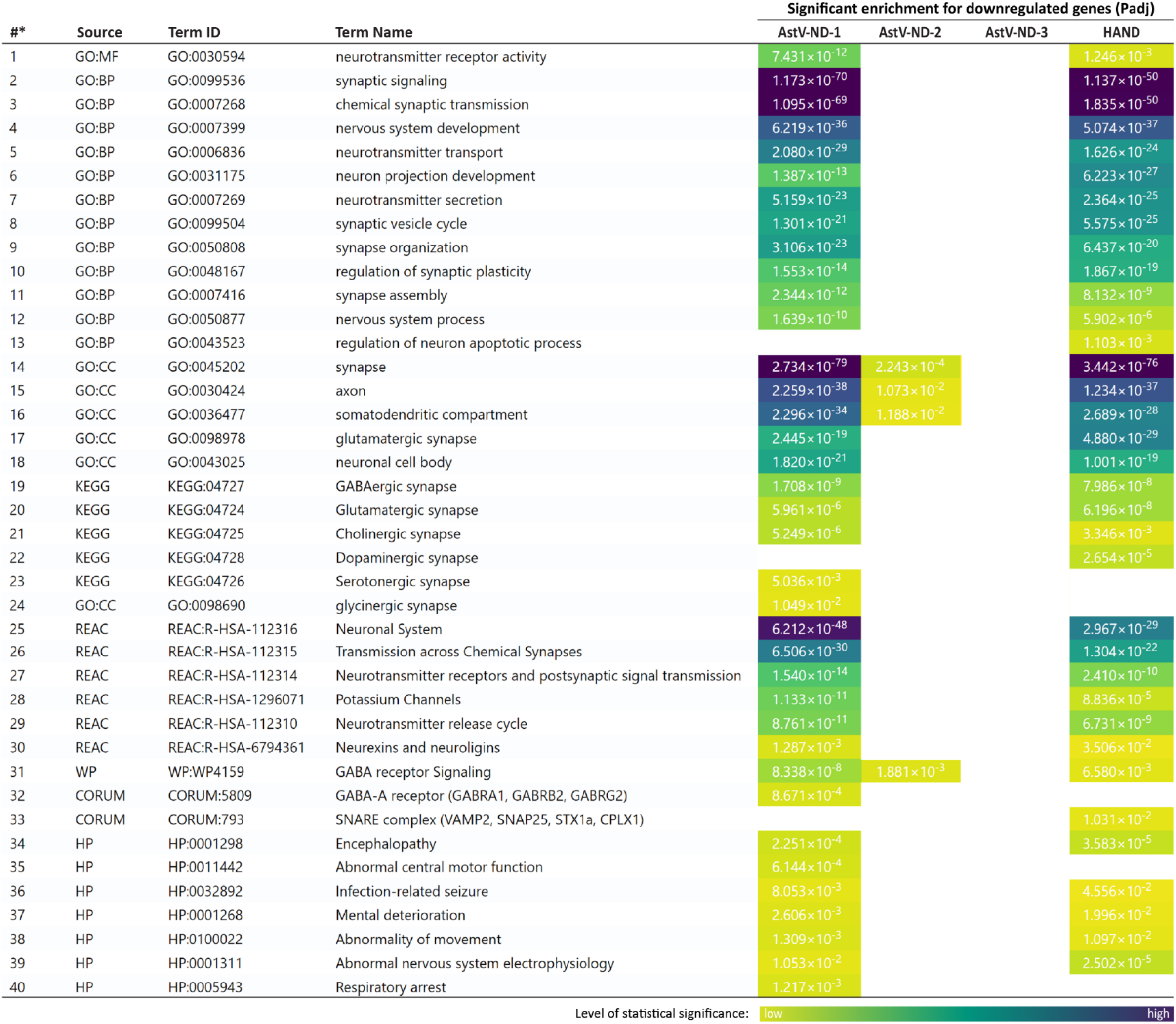
Multi-source side-by-side heatmap of the statistical significance for functional enrichment related to the downregulated gene expression in AstV-ND and HAND. Multi-source functional enrichment terms of interest and their corresponding Padj enrichment values for indicated cases of astrovirus and HIV CNS infections. Blanks indicate the absence of significance (Padj > 0.05). *Indicates the term enumeration corresponding to that shown in in Fig. 5 (solid color circles). Enrichment values are colored according to the level of statistical significance (scale is provided at the bottom of the figure).

Strikingly, the AstV-ND-1 was very similar to HAND with respect to the magnitude and repertoire of transcriptional downregulation, with enrichment of virtually the same functional terms that signified impairment of neural functions (Figs. 5 and 6). The most significantly enriched terms in both AstV-ND-1 and HAND were “synaptic signaling” (term #2); “chemical synaptic transmission” (term #3); “synapse” (term #14); and “neuronal system” (term #25). Both AstV-ND-1 and HAND also had similar significant enrichments in human disease associations which included the HP terms “Encephalopathy” (term # 34); “Infection-related seizure” (term #36); “Mental deterioration” (term #37); “Abnormality of movement” (term #38); and “Abnormal nervous system electrophysiology” (term #39). AstV-ND-1 and HAND also had a largest overlap (n = 124) in specific downregulated genes (Fig. 7a; all genes associated with the Venn diagram are listed in Supplemental File 7). Analysis of these downregulated genes revealed a common functional signature indicative of dysregulation of neurotransmission and impairment of neural functions associated with physiological changes in the synapses, axons, and somatodendritic compartments of neurons (Fig. 7, b and c; complete list of significantly enriched terms corresponding to Fig. 7b is provided in Supplementary 8).

**Figure 7.**
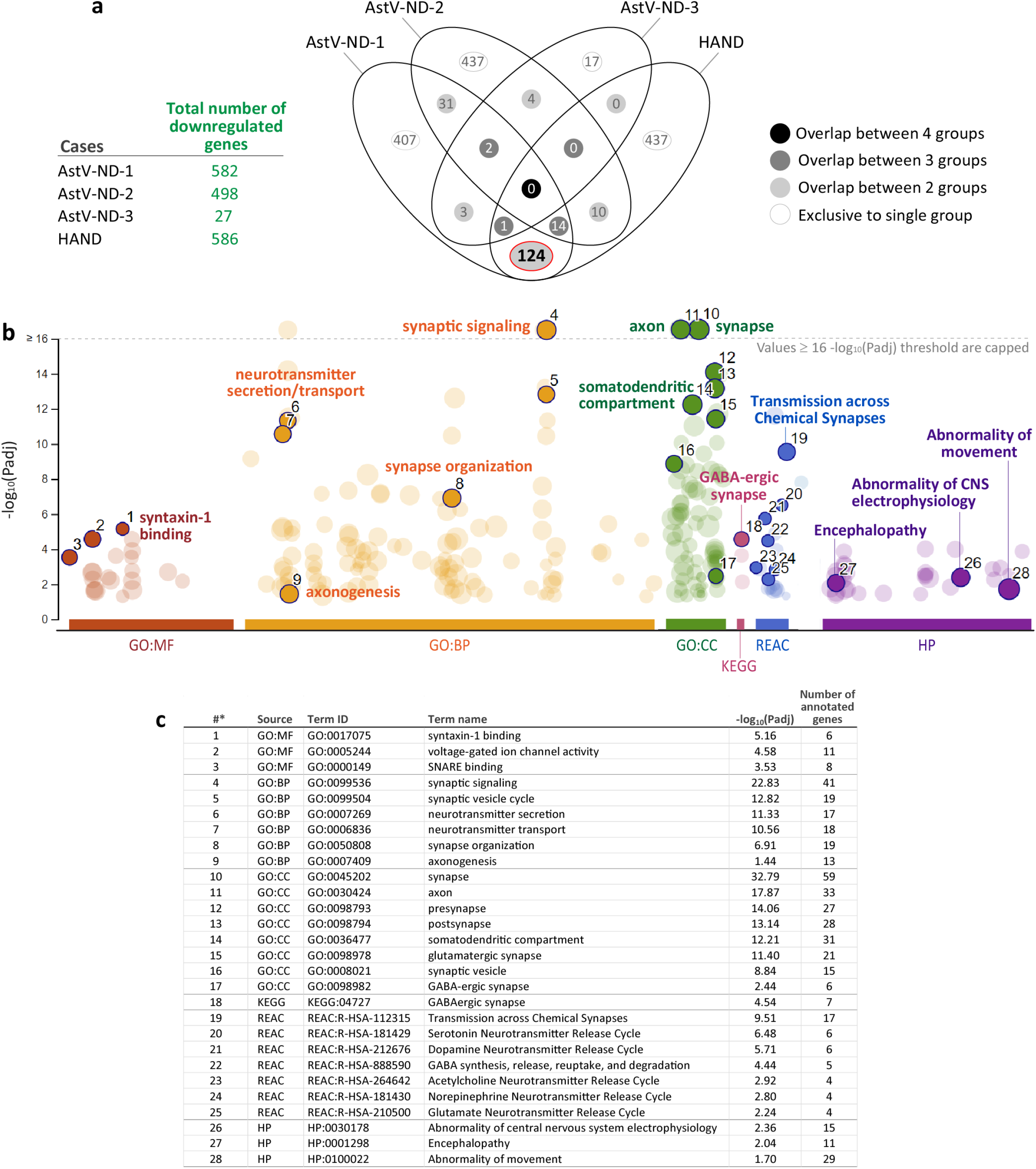
Functional genomic analysis of the common downregulated genes in AstV-ND and HAND. (**a**) Venn diagram comparison of the groups of genes that were downregulated in indicated cases of astrovirus and HIV CNS infections. (**b**) Manhattan plot shows significantly enriched terms (Padj < 0.05) across multiple ontologies for genes that were commonly downregulated in the largest overlap (highlighted by red border in **a**). Plot features and gene ontology data source abbreviations are as in Fig 2. (**c**) Statistical details for the enriched terms enumerated (denoted as #*) and shown by solid circles in **b**.

Since brain tissue from AstV-ND-1 had a much larger number of downregulated genes compared to other two cases of AstV-ND, we analyzed the downregulated neural functions in AstV-ND-1 case by employing an ordered query in gProfiler (https://biit.cs.ut.ee/gprofiler/page/docs#ordered_gene_lists) which allows differential gene expression to be ranked and placed in a biologically meaningful order from higher to lower importance. This analysis placed downregulated functions associated with the synapse, axon, and somatodendritic compartment at the top of the list (Fig. 8a). Functional terms with major “spikes” in the level of statistical significance for downregulated genes included: synapse (138 downregulated synaptic genes are listed in Fig. 8b), synaptic signaling, presynapse, synaptic membrane, synaptic vesicle cycle, and neurotransmitter secretion. The most affected synaptic types were the glutamatergic and GABA-ergic synapses. Terms such as synapse organization, regulation of synaptic plasticity, synapse assembly, protein-protein interactions at synapses, and neurexins and neuroligins were also among the top-ranked downregulated functions (complete list of significantly enriched terms corresponding to Fig. 8a is provided in Supplementary File 9).

**Figure 8.**
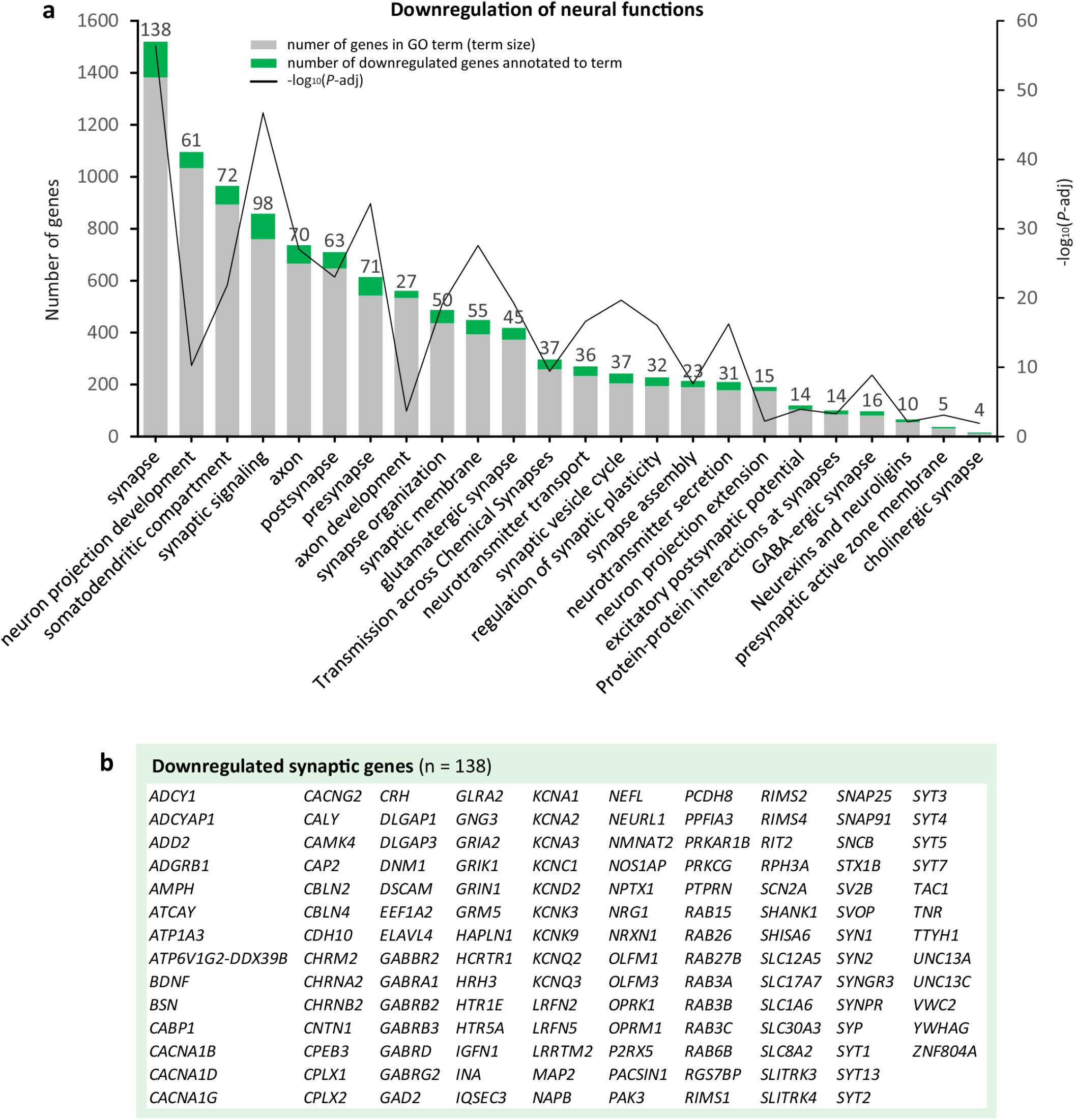
Computational dissection of the downregulated neural functions in AstV-ND-1. (**a**) Selected significantly enriched (FDR < 0.05) gene ontology (GO) terms (Biological Process and Cellular Component) for genes that were downregulated in the brain of AstV-ND-1 as compared to all normal brain control samples (ordered gProfiler query for genes downregulated in AstV-ND-1 vs. all normal controls; see details for normal controls in Fig. 1, a and c). Significance is shown on the y axis as -log_10_ of the adjusted P value. (**b**) List of significantly (FDR < 0.05; fold change < 2.0) downregulated genes (n = 138) associated with the GO term “synapse”.

To get further insight into upstream regulation this neural impairment signature, we used the TRANSFAC database via g:GOSt prediction pipeline and identified putative transcription factors (TFs) (Supplemental Fig. 2). The most strongly implicated motifs were associated with the neuron-restrictive silencing transcription factor RE1-Silencing Transcription factor (*REST*), also known as Neuron-Restrictive Silencer Factor (*NRSF*). *REST/NRSF* encodes a protein which represses transcription of neuron-specific genes by binding the neuron-restrictive silencer element (*NRSE*, also known as *RE1*) and may act as a master transcriptional regulator in neurodegenerative diseases [19].

Together, these findings showed a high similarity in extensive transcriptional downregulation and impairment of neural functions associated with opportunistic CNS infections caused by astrovirus in an adult with acquired iatrogenic immunodeficiency (AstV-ND-1, 58-years of age) and adults with HAND (with acquired HIV-1-associated immunodeficiency, 43 to 58 years of age). In contrast, AstV-ND in children with a primary immunodeficiency (XLA, 14-years of age [AstV-ND-3] and 15-years of age [AstV-ND-2]) was associated with modest downregulation of neural functions in a case of a fatal encephalitis (AstV-ND-2) and minimal downregulation of neural functions in a patient who survived the infection (AstV-ND-3).

**Supplemental Figure 2.**
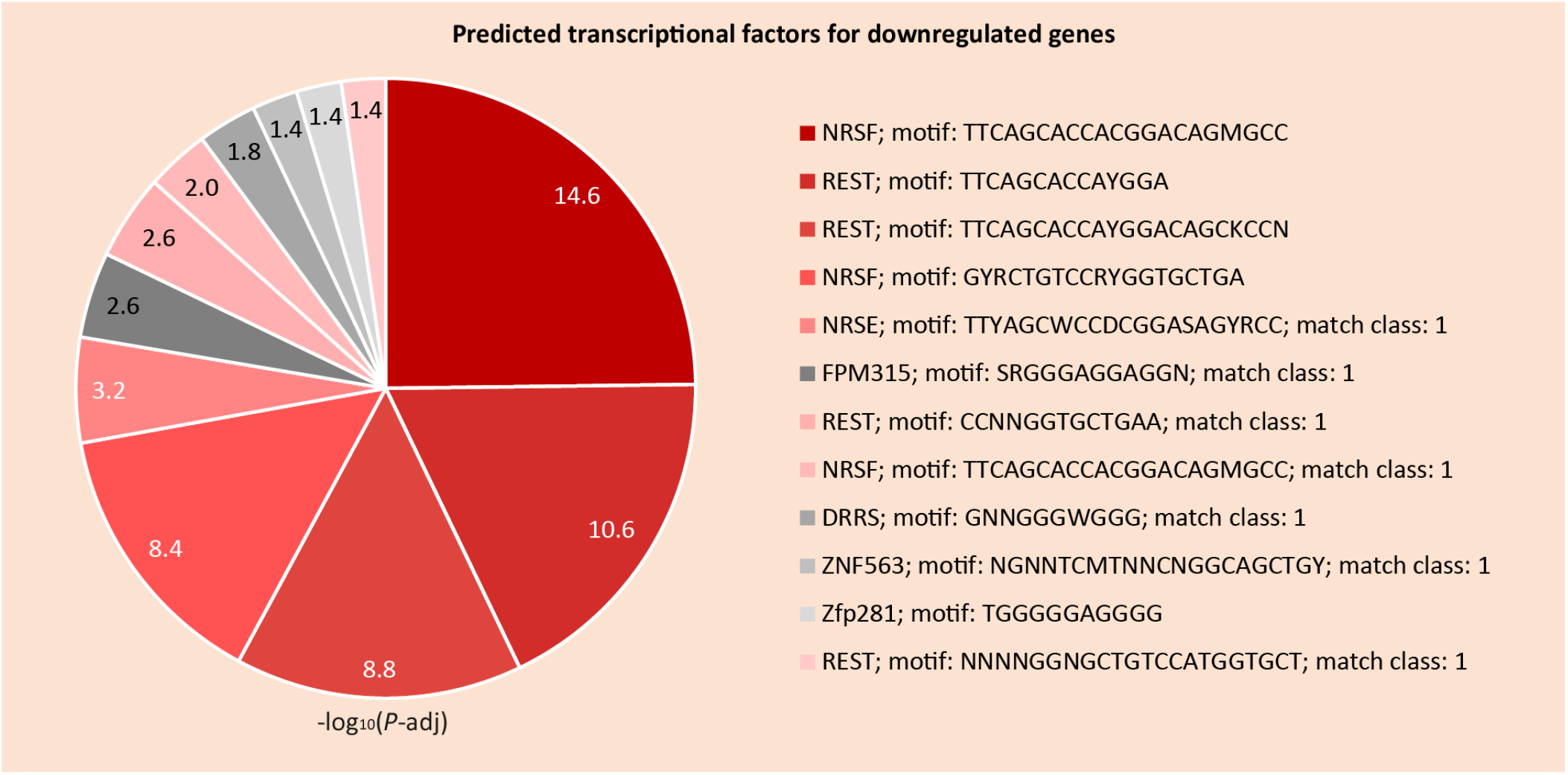
Computational prediction of potential transcription factors (TFs) for genes downregulated in the brain of patient AstV-ND-1. The relative significance depicted in the pie chart is based on -log_10_ FDR-adjusted p values; corresponding motifs/match classes are shown on the right. We used the TRANSFAC database via g:GOSt prediction pipeline (https://biit.cs.ut.ee/gprofiler/page/docs#ordered_gene_lists) that minimizes the number of false positive matches (provided by TRANSFAC) to remove spurious motifs. Remaining matches were split into two groups based on the number of matches: (i) base motif class that contained all genes matching a given motif and (ii) motifs that had at least two matches per gene (classified as match class 1).

### Human astrovirus targets neurons, especially in the brainstem, and disseminates along neuronal projections

Brain tissue for immunohistochemistry was only available from AstV-ND-1; therefore, we probed these tissues with antibodies to both astrovirus and cellular proteins. Both glial cells (e.g., astrocytes) [6] and neurons [6, 9, 10] have been reported to contain astrovirus RNA or capsid protein in patients with astrovirus encephalitis. Therefore, we used immunohistochemistry with a well-established antibody against astrovirus capsid protein [6, 10] and analyzed all major CNS regions to identify the type of cells that were infected. The CNS regions analyzed were cerebral cortex (frontal and temporal), hippocampus, basal ganglia, thalamus, cerebellum (cerebellar cortex and deep cerebellar nuclei), and brainstem (i.e., midbrain [substantia nigra], pons, and medulla oblongata [including inferior olives]).

All these CNS regions harbored astrovirus capsid protein in only neurons (Fig. 9, a - h). Virus capsid protein was found not only in the perikarya of neurons, but also in their projections (dendrites and axons) over a substantial distance from the neuronal body. The topology and morphology of astrovirus infected cells, as well as additional double immunofluorescent staining for astrovirus capsid protein and microtubule associated protein 1 (MAP2; pan-neuronal somatodendritic marker) (Fig. 9, i - k), unambiguously identified neurons as the cell type supporting astrovirus infection.

**Figure 9.**
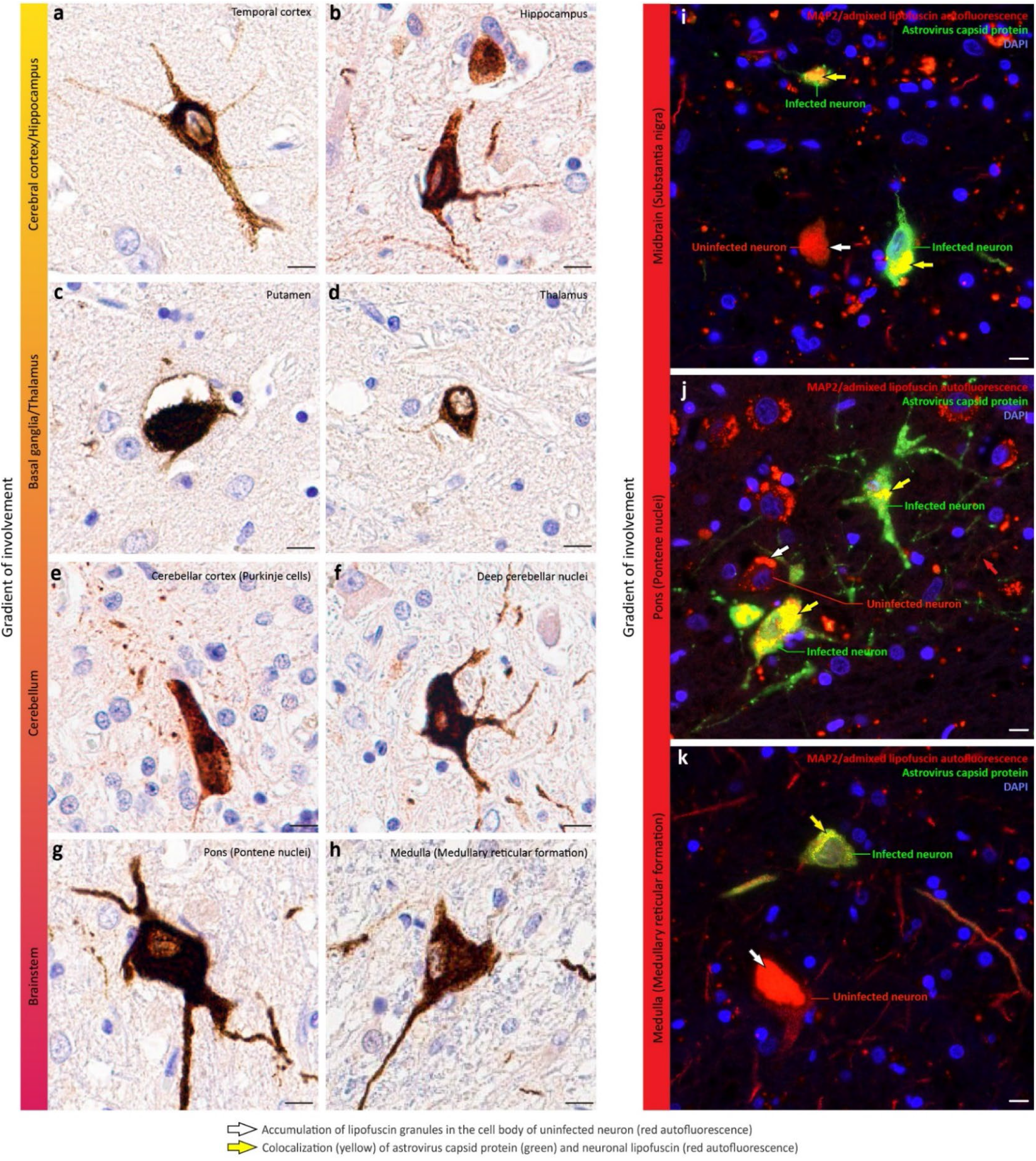
Gradient of involvement of the neurons along the neural axis in astrovirus infection. (**a - h**) Astrovirus capsid protein (brown) in neuronal perikarya and projections (dendrites and axons) is present in various CNS structures. Gradient of involvement (from yellow [restricted] to red [extensive]) of indicated CNS regions by astrovirus infection of neurons is shown on the left. (**i** - **k**) Double immunofluorescent staining for the neuronal somatodendritic marker MAP2 (red), astrovirus capsid protein (green), and DAPI nuclear counterstain (blue) in the indicated CNS structures with extensive involvement. Note: accumulation of aging pigment lipofuscin, which is known to be autofluorescent, produces an intense autofluorescent signal localized to the neuronal cell bodies in uninfected neurons (red; MAP2 with admixed autofluorescence) and this signal colocalizes with the specific immunofluorescent signal for the astrovirus capsid protein (green) in astrovirus-infected neurons (**i** - **k**). Labeling keys used in (i - k) are provided at the bottom of the figure. Scale bars: 10 μm.

Only a few infected neurons were readily detected in the cerebral cortex, hippocampus, basal ganglia, and thalamus (Fig. 9, a - d). The cerebellum harbored more numerous infected neurons (i.e., inhibitory Purkinje cells in the cerebellar cortex and large excitatory neurons in the deep cerebellar nuclei; Fig. 9, e and f). Of all the areas of the brain examined, the brainstem (i.e., midbrain, pons, and medulla oblongata) harbored the most numerous astrovirus infected neurons (Fig. 9, g - k). Thus, neurons are the major cell type capable of supporting astrovirus replication in human CNS and the brainstem followed by the cerebellum had the highest numbers of virus infected neurons.

### The cellular response to human astrovirus infection in the CNS is dominated by intrinsic and extrinsic phagocytic cells

Next, we used immunohistochemistry to analyze CNS regions that harbored astrovirus-infected neurons in AstV-ND-1. Scattered mild perivascular inflammatory cell infiltrates and perineuronal hypercellularity were observed, especially in the brainstem (Fig. 10, a and b; H&E). Many CNS regions showed robust responses by phagocytic cells with CD68 immunoreactivity which were most prominent in the brainstem. These included many CD68+ perivascular phagocytic cells (presumably including the resident perivascular macrophages and peripheral extravasated macrophages), some of which appeared to infiltrate the surrounding brain parenchyma (Fig. 10c). CD68+ cells with a morphology consistent with activated microglia surrounded many brainstem neurons (Fig. 10d). A moderate astrocytosis in the brainstem was characterized by the hypertrophy of astrocytic somata, perivascular end foot processes (Fig. 10e), and perineuronal processes (Fig. 10f). Hypertrophic perineuronal astrocytic processes appeared to retract from the somata of neurons, which could allow other inflammatory cells (e.g., activated microglia/macrophages; compare Fig. 10f to Fig. 10d) to localize in close apposition to neurons. In contrast, there was a very limited infiltration by CD3+ T cells (Fig. 10g) and these lymphocytes were not observed adjacent to neurons. As expected, given the B-cell depleting therapy received by the patient, perivascular infiltrates did not contain CD20+ B cells (Fig. 10h).

**Figure 10.**
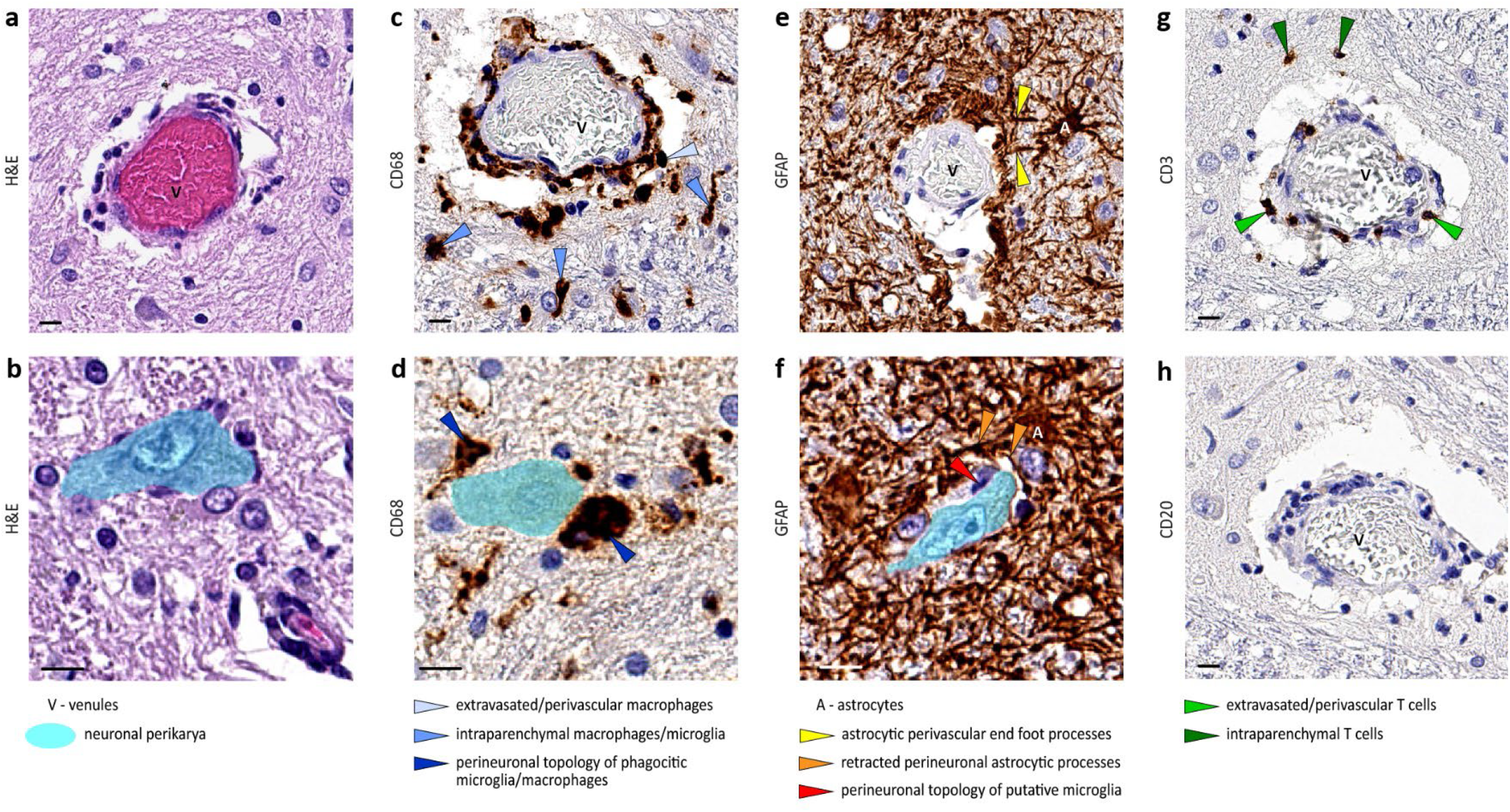
Cellular response to astrovirus infection in the brainstem. **(a - h)** Representative images show the perivascular (a, c, e, g, and h) and perineuronal (b, d, and f) tissue in adjacent sections of the medulla that illustrate: (1) perivascular and perineuronal hypercellularity (Hematoxylin and eosin [H&E]; a and b); (2) perivascular and perineuronal topology of activated macrophages/microglia (brown; CD68; c and d); (3) moderate astrocytosis (brown; glial fibrillary acidic protein [GFAP]; e and f) with (i) hypertrophy of astrocytic somata and perivascular end foot processes (e) and (ii) retraction of perineuronal astrocytic processes and a cell with microglial morphology that is in close apposition to the neuronal membrane; (4) infiltrating T cells (brown; CD3; g); and (5) absence of infiltration by B cells (CD20; h). Keys for labeling are provided below each respective pair of panels. Scale bars: 10 μm.

Together, these findings show that the cellular response to astrovirus infection in the CNS was dominated by activated microglial cells (intrinsic glial cells with an activated phagocytic phenotype) and infiltrating peripheral macrophages, indicating ongoing phagocytosis. Astrocytic responses were moderate and appeared to act in limiting parenchymal infiltration by peripheral immune cells from perivascular spaces by hypertrophy of their perivascular end feet processes, while allowing access for activated microglia/macrophages to infected neurons by retraction of their perineuronal processes. The brainstem, which harbored most astrovirus-infected neurons, showed the most intense cellular response to the infection.

### Human astrovirus infection disrupts synapses and impairs both excitatory and inhibitory neurotransmission

As noted above many synaptic genes were downregulated in astrovirus-infected brain (Fig. 8b) indicating impairment of neural functions. Therefore, we next used immunohistochemistry to interrogate this impairment at the protein expression level. We selected three significantly downregulated synaptic genes that were functionally annotated to the (1) synaptic vesicles (synaptophysin [*SYP*]; pan-presynaptic marker), (2) excitatory glutamatergic synapses (solute carrier family 17 member 7 [*SLC17A7*]; encoding vesicular glutamate transporter 1 [VGLUT1]; presynaptic marker of glutamatergic synapses), and (3) inhibitory synapses (solute carrier family 32 member 1 [*SLC32A1*]; encoding vesicular GABA transporter [VGAT]; presynaptic marker of GABA-ergic synapses) (Supplemental Fig. 3).

Immunohistochemical staining of the thalamus, midbrain, pons, and medulla samples from the patient with AstV-ND-1 using a pan-presynaptic marker synaptophysin showed a loss of the presynaptic terminals in the neuropil surrounding astrovirus-infected neurons (Fig. 11) This suggested that astrovirus replication in neurons triggers the loss of their afferent innervation and leads to impaired neurotransmission. Consistent with this, there was a marked reduction of both excitatory (VGLUT1+) and inhibitory (VGAT+) presynaptic puncta in the brainstem of the patient with AstV-ND-1 (compared to a normal age-matched control) (Fig. 12).

**Figure 11.**
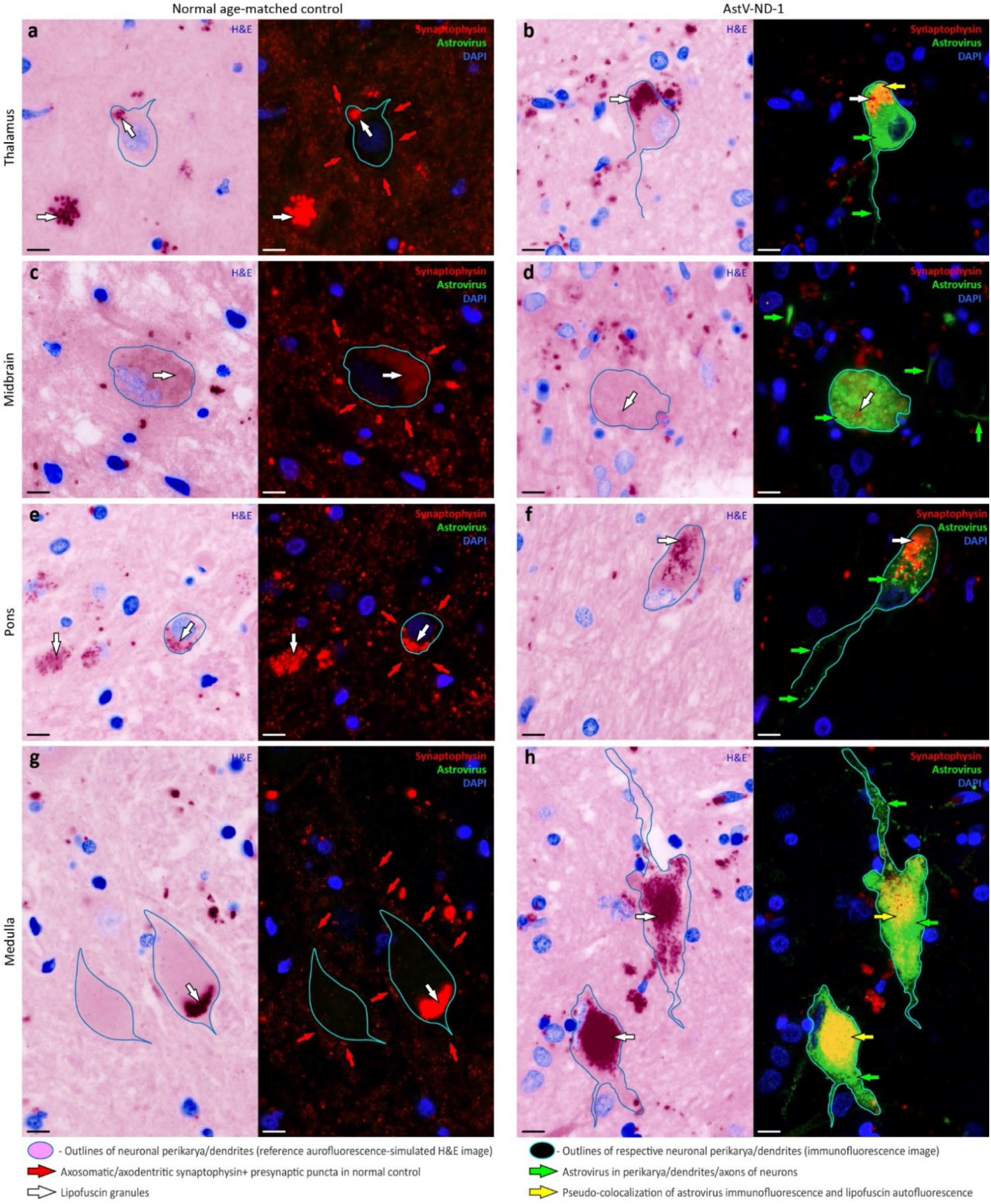
Astrovirus replication in neurons triggers loss of their afferent innervation. (**a** - **h**) Representative images show a side-by-side comparison of the density of synaptophysin-immunoreactive presynaptic terminals in the neuropil surrounding uninfected neurons (“Normal age-matched control” column) versus astrovirus-infected neurons (“AstV-ND-1” column) in indicated brain regions. Each panel is composed of (1) an autofluorescence-simulated image that was pseudo-colored as hematoxylin-eosin (H&E) staining to serve as a topographical reference and to aid identification of intra- and extra-neuronal autofluorescent lipofuscin granules in corresponding immunofluorescent images (see Materials and Methods) and (2) immunoreactivity signals for the pan-synaptic marker synaptophysin (red), astrovirus (green), and DAPI nuclear counterstain (blue) in the corresponding tissue field. Outlines of the select neuronal perikarya/dendrites correspond to the same neurons shown in the H&E reference images and corresponding immunofluorescent images. Labeling keys used in (a - h) are provided at the bottom of the figure. Scale bars: 10 μm.

**Figure 12.**
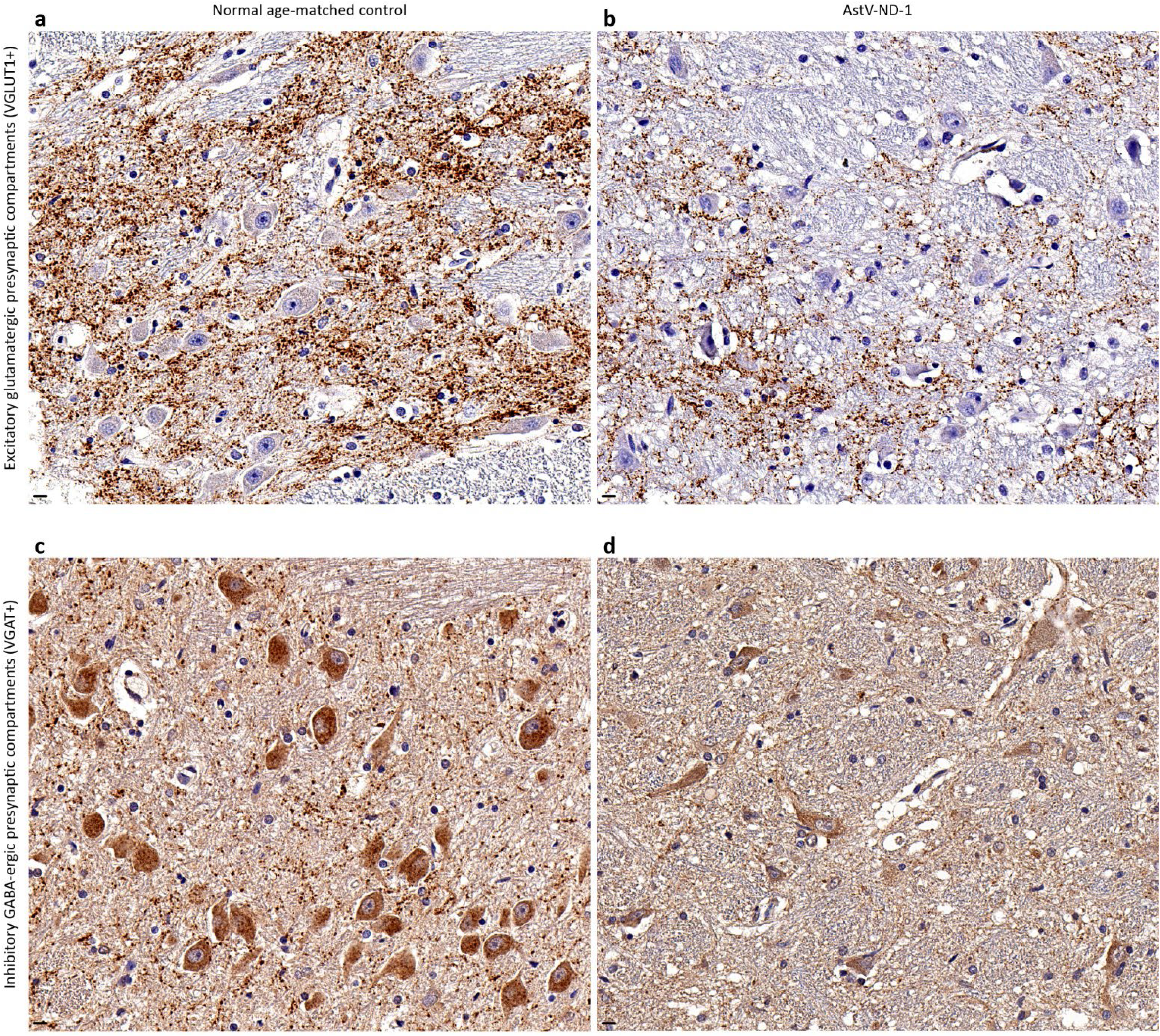
Astrovirus infection is associated with impairment of both excitatory and inhibitory neurotransmission. (**a** - **d**) Representative images show a side-by-side comparison of the density (brown immunoreactivity) of presynaptic compartments of excitatory glutamatergic VGLUT1+ synapses (a and b) and inhibitory GABA-ergic VGAT+ synapses (c and d) in the brainstem of a normal age-matched control subject (a and c) and in the brainstem of patient with AstV-ND-1 (b and d). Note a loss of the excitatory and inhibitory presynaptic puncta in the brainstem of patient with AstV-ND-1. Scale bars: 10 μm.

Taken together, these findings demonstrate that astrovirus infection in the human brain disrupts the synaptic integrity, triggers the loss of afferent innervation related to infected neurons, and leads to a global impairment of both excitatory and inhibitory neurotransmission.

**Supplemental Figure 3.**
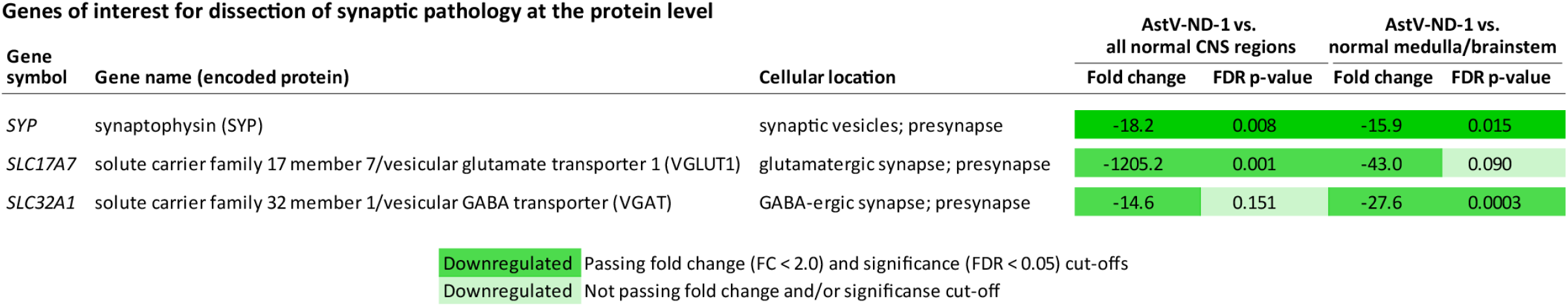
Details on expression of three significantly (chosen cut-offs: FDR < 0.05; fold change [FC] < 2.0) downregulated genes of interest selected based on data shown in Fig. 8, a and b. The FCs and FDR p-values were different when comparing AstV-ND-1 to all normal CNS regions (since the site of brain for the sample from AstV-ND-1 was unknown), or to only the normal medulla/brainstem (putative site of AstV-ND brain sample based on its significant association with human disease phenotype terms related to functions of the brainstem [“Abnormal central motor function” (term #35) and “Respiratory arrest” (term #40)]; Fig. 6).

## DISCUSSION

Astroviruses are enteric RNA viruses [4] that occasionally result in CNS infections in humans, mainly in patients with congenital (primary) immunodeficiencies or hematologic malignancies undergoing immunosuppressive treatments [6-14] (Table 1). While some viruses invade the CNS shortly after primary infection (e.g., arboviruses such as West Nile virus), other viruses infect the CNS after reactivating from latent infection that was established many years in the past (e.g., herpes simplex, varicella-zoster virus, and JC virus). Such latent viruses while controlled in immunocompetent individuals, may invade the CNS when presented an “opportunity” due to impaired immune control in immunocompromised individuals. Astrovirus neurological disease in patients with a primary or secondary immunodeficiencies (summarized in Table 1) represents an opportunistic CNS infection. Here we identified a novel human neuropathogenic astrovirus (HAstV-NIH) that appeared to be gastrointestinal in origin (Supplemental Figure 1) and closely related to other human astroviruses that were detected in the CNS of immunocompromised patients and to bovine astroviruses linked to encephalitis in cattle.

We unambiguously demonstrate that CNS neurons are target cells infected by astrovirus. This is consistent with most reports where astrovirus infection (astroviral RNA or capsid protein) was detected in neurons of infected humans or animals (Table 1). We show that astrovirus capsid protein is present not only in the neuronal perikarya, but also in their dendrites and axons over a substantial distance from the neuronal body. This strongly suggests that astrovirus can spread along neuronal projections. Intriguingly, we found astrovirus capsid protein in the same brain structures (i.e., thalamus, deep cerebellar nuclei, Purkinje cells, and pontine nuclei) that are preferentially infected in primates by West Nile virus [20]. Based on the neuroanatomy and known neural connectivity, we postulate that following invasion of the CNS, astrovirus spreads trans-synaptically in a retrograde manner (e.g., thalamus -> deep cerebellar nuclei -> Purkinje cells) between connected neurons. Therefore, as has been shown for West Nile virus infection [21], transsynaptic propagation of astrovirus infection within the CNS may result in profound neurophysiological impairment. It appears that the gradient of neuronal astrovirus infection in the human brain intensifies caudally with the major impact on the brainstem, followed by the cerebellum. Interestingly, a similar caudal gradient of neuronal involvement is present in the CNS of astrovirus-infected cattle (summarized in Table 1).

The human CNS has barriers to protect itself from an invasion by various pathogens (including viruses) that otherwise would be detrimental to neural function and could be life-threatening. Viremia ensues when virus replication in the peripheral tissue reaches a threshold and the virus enters the bloodstream. Viremia facilitates virus spread to various tissues (including peripheral nerves) and increases the probability of virus invasion of the CNS. Indeed, viremia is a major neuropathogenic determinant for CNS infection [22]. However, immunocompetent hosts with intact innate and adaptive immunity are usually able to contain virus replication in the periphery, reduce the level of viremia, and prevent virus invasion of the CNS. In contrast, immunocompromised persons who fail to mount successful innate and/or adaptive antiviral immune responses in the periphery may develop devastating neurological disorders associated with viral neuroinvasion. These persons may have primary immunodeficiencies (such as genetic disorders that impair the innate and/or adaptive immune response), secondary immunodeficiencies (including recipients of immune-cell depleting and/or immunomodulatory therapies or those with AIDS), or being persons at the extremes of age (including infants with immature immune systems or the elderly with immunosenescence) [2].

Underlying conditions in patients with AstV-ND in this study included the primary and secondary immunodeficiencies (Table 1) predominantly affecting B cell immunity (XLA or lymphoma/leukemia with B-cell depleting and immunosuppressive treatments). Adaptive immunity, including B cell activity, is important to contain viral infections in the periphery to prevent invasion of the CNS, as well as to promote cytolytic and non-cytolytic virus clearance in the CNS (reviewed in [23, 24]). Our findings show that defective adaptive immunity in patients with AstV-ND impairs effective responses to astrovirus infection within the CNS, since gene expression in the brain did not exhibit significant changes indicative of activation of adaptive immunity. Nevertheless, we detected robust changes in gene expression in the brain associated with activation of innate immune responses to astrovirus infection. This phenomenon was further corroborated by functional patterns of gene upregulation in the brain and immunohistochemistry of CNS tissues, all of which was dominated by activation of microglia and macrophages. As expected, deficiency in B cells whether due to XLA [25] or hematologic malignancies with B-cell depleting and immunosuppressive treatments, results in the absence of B cell/plasma cell migration to the brain to fight astrovirus infection. Indeed, similar to reported cases of the CNS astrovirus infections in immunocompromised persons (summarized in Table 1), we found only minimal lymphocytic infiltration of the brain which was composed predominantly of T cells and peripheral macrophages. In contrast, astrovirus infection of the CNS in animals is often associated with prominent perivascular and leptomeningeal lymphocytic infiltration (Table 1) with admixed plasma cells [26]. Interestingly, astrocytic retraction of their perineuronal processes appeared to play a supportive role by facilitating access for activated microglia/macrophages to position themselves close to infected neurons. On the other hand, hypertrophy of perivascular astrocytic end feet may limit excessive parenchymal infiltration by peripheral immune cells from perivascular spaces. Taken together, these findings suggest that innate immune responses and phagocytosis have an important role in the neuropathogenesis of opportunistic AstV-ND in patients with underlying deficiencies in adaptive immunity.

Astrovirus infection of the CNS in immunocompromised patients is often fatal and the brainstem is usually the most severely affected structure (Table 1). Our findings expand understanding of functional impact of astrovirus infection on neurophysiology. We show that astrovirus infection leads to physiological changes in neurons that include transcriptional downregulation of genes associated with the neuronal somatodendritic compartments, axons, as well as the excitatory and inhibitory synapses. Immunohistochemistry confirmed a loss of the presynaptic terminals in the neuropil surrounding astrovirus-infected neurons and a corresponding loss of vesicular transporters associated with both excitatory and inhibitory synapses. This shows that astrovirus infection of neurons impacts neurophysiology by disrupting synaptic integrity, triggering a loss of afferent innervation related to infected neurons, and impairing both excitatory and inhibitory neurotransmission. To date, relatively few studies have investigated the downregulation of expression of neural genes [17, 27] or synaptic degeneration and loss [28-32] in postmortem brains of humans who died from CNS infections caused by viruses (e.g., HIV and WNV). However, there are many examples of downregulation of neural functions at the gene and protein level, as well as synaptic pathology, in animal studies of CNS infections caused by various viruses [21, 33-35].

The CNS defense response to astrovirus infection in immunocompromised patients shares many features with other opportunistic viral infections. For instance, we found shared functional patterns of gene upregulation in the brain in response to opportunistic virus CNS infections (AstV-ND and HAND) that were dominated by innate immune response with microglial and macrophagic signatures whereas regulation of adaptive responses was impaired. This may be a common compensatory defense mechanism employed by the CNS against opportunistic infections in humans with impaired adaptive immunity. With the increasing use of immunosuppression for a variety of disorders (e.g., autoimmune disease, organ/stem cell transplantation, and cancer) predisposing to opportunistic viral infections of the CNS, as well as the emergence or reemergence of neurotropic viruses (e.g., enterovirus D68, West Nile, Chikungunya, Zika, Hendra, and Nipah viruses) [3, 36] the frequency of viral encephalitides is expected to increase in the coming years. In addition, global warming may contribute to spread of viral vectors in new areas that can increase the prevalence of arthropod borne viruses to new populations [37]. These changes will continue to present major challenges for the diagnosis and treatment of these diseases as well as for public health measures to limit the spread of these viruses. Since effective treatments do not exist for most opportunistic viral infections of the CNS, the only option is the reversal of immunosuppression, with the risk of worsening the underlying condition [36]. Thus, a better understanding of the pathophysiology of opportunistic viral infections, including astrovirus encephalitis, is needed to limit neurologic damage associated with these diseases and to develop new therapeutic approaches.

## MATERIALS AND METHODS

### Brain tissue samples

For RNA-seq, frozen brain tissue samples from three patients with AstV-ND were used: (i) AstV-ND-1 (unknown brain site; 58-year-old man; see case description below); (ii) AstV-ND-2 (brainstem; a 15-year-old boy with X-linked agammaglobulinemia [XLA]) [6]; and (iii) AstV-ND-3 (frontal cortex; a 14-year-old boy with XLA) [8]. Frozen brain tissue samples from subjects (ages 14 to 58-years old) without known neurological disease were used as normal controls (frontal cortex [n=2]; thalamus (n=1); brainstem [n=1]; medulla [n=1]; and cerebellum [n=1]). These normal tissue samples were obtained from the Human Brain Collection Core, Intramural Research Program, NIMH (http://www.nimh.nih.gov/hbcc). Formalin-fixed paraffin-embedded (FFPE) brain tissue sections from AstV-ND-1 case were used for histopathological analysis (hematoxylin-eosin [H&E] staining), brightfield colorimetric immunohistochemistry, and double immunofluorescent staining. FFPE sections from an age-matched subject without known neurological disease were used as normal controls.

Consent for research studies was obtained for use of leftover brain tissue from AstV-ND-3 from the patient’s parents; brain tissue was obtained postmortem from AstV-ND-1 and AstV-ND-2 and as such is not considered human subjects research.

### AstV-ND-1 case description

A 58-year-old man presented with altered mental status after receiving a double unit umbilical cord blood transplant for lymphoma. The patient was diagnosed with follicular lymphoma, was treated with rituximab, cyclophosphamide, doxorubicin, vincristine, and prednisone therapy for 6 cycles and underwent remission. He was diagnosed with recurrent disease three years later and was treated with radiation therapy. Four months later biopsy of a new soft tissue mass showed diffuse large B cell lymphoma compatible with transformation of follicular lymphoma, and he received three cycles of rituximab, dexamethasone, cytarabine, and cisplatin therapy. He received conditioning with cyclophosphamide, thiotepa, fludarabine, and total body irradiation (400 cGy) and underwent a double cord blood stem cell transplant. He received cyclosporine and mycophenolate mofetil for graft-versus-host disease prophylaxis. He engrafted with 100% donor cells but developed gastrointestinal graft-versus-host disease and was treated with methylprednisolone and budesonide and his cyclosporine and mycophenolate mofetil were continued. One month after transplant, the patient presented with lower extremity sensory changes and weakness with ascending neuropathy, ataxia, unstable gait, and fatigue. An MRI of the brain and cervical spine were unremarkable, and an EMG showed a sensory motor neuropathy. The cerebrospinal fluid (CSF) showed 5 white blood cells/ml, protein 55 mg/dL, glucose 57 mg/dL and was negative for human herpesvirus 6, varicella-zoster virus, herpes simplex virus, JC virus, and cytomegalovirus. Cytology showed atypical lymphocytes. A paraneoplastic panel was negative and oligoclonal bands were not detected. Three months after his neurologic changes began, a PET scan of brain showed a mildly abnormal thalamus with increase uptake, and one week later, an MRI showed a new fluid attenuated inversion recovery hyperintensity with restricted diffusion in the right midbrain, both cerebellar peduncles, and right lateral pons. An EEG showed background slowing, and the CSF showed protein 48 mg/dL, glucose 65 mg/dL, white blood cell count 5 cells/ml with 68% lymphocytes, and many atypical lymphocytes and was again with negative for human herpesvirus 6, JC virus, and cytomegalovirus. He was treated with acyclovir and foscarnet followed by rituximab and solumedrol. He developed progressive mental status changes, weakness, extra-pyramidal signs, and agitation. An EEG showed generalized slowing and an EMG showed a sensorimotor polyneuropathy involving the bilateral lower extremities that was ascending. CSF showed elevated protein and lack of pleocytosis, and he was treated with high dose intravenous immunoglobulin and steroids for presumptive variant Guillain-Barre. A sural nerve biopsy showed nerve damage. He received an additional course of rituximab and died seven months after transplant. At autopsy, the diagnosis was a multifocal leukoencephalopathy of unknown etiology.

### Gene expression analysis

Tissue samples were homogenized for 40 sec in lysing matrix D tubes (MP Biomedicals, Santa Ana, CA) containing 1 ml Trizol (Thermofisher Scientific, Waltham, MA) in a FastPrep® FP 120 instrument (MP Biomedicals) at a speed 6.0 meters per second. Homogenized Trizol lysate was combined 1-Bromo-3-chloropropane (MilliporeSigma, St. Louis, MO), mixed, and centrifuged at 16,000 x g for 15 min at 4°C. RNA containing aqueous phase was collected from each sample and passed through a Qiashredder column (Qiagen, Valencia, CA) at 21,000 x g for 2 min to homogenize any remaining genomic DNA in the aqueous phase. The aqueous phase was combined with an equal amount of RLT lysis buffer (Qiagen, Valencia, CA) with 1% beta-mercaptoethanol (MilliporeSigma, St. Louis, MO), and RNA was extracted using Qiagen AllPrep DNA/RNA mini columns (Valencia, CA). The RNA yield ranged from 860 ng to 12.4 μg. RNA quality was determined by spectrophotometry at 260 nm and 280 nm and by fluorescence capillary assay (RNA 6000 Pico kit, Agilent Technologies, Santa Clara, Ca). RNA integrity numbers ranged from 2.3 to 6.2.

A Truseq Stranded mRNA-Seq Sample Preparation Kit (Illumina) was used to synthesize cDNA and generate sequencing ready libraries following the manufacturer’s protocol. Due to RNA degradation and low RNA integrity values, the fragmentation time was reduced from eight to six min. Library quality and size distribution was assessed on a BioAnalyzer DNA 1000 chip (Agilent Technologies), and concentrations were determined using the Kapa SYBR FAST Universal qPCR kit for Illumina sequencing (Roche). Paired-end 75 cycle sequencing was completed using two Mid Output 150 cycle kits on the NextSeq 550 (Illumina).

Raw reads were trimmed of adapter sequence using cutadapt (https://cutadapt.readthedocs.io/en/stable/). The remaining reads were then filtered for low quality bases and low quality reads using the FASTX-Toolkit (http://hannonlab.cshl.edu/fastx_toolkit/). The remaining reads were mapped to the GRCh38 genome, using HISAT2. Reads mapping to genes were counted using htseq-count.

Differential expression analysis was performed using the Bioconductor package DESeq2 along with generation of a sample heatmap using the pheatmap package with default parameters (Supplemental figure 5).

Clustering analysis of the transcriptomes from the three tissues from patients with AstV-ND with brain tissues from normal controls (thalamus, brainstem, medulla [part of brainstem], frontal cortex, and cerebellum) showed that AstV-ND-2 [brainstem] transcripts clustered with brainstem and medulla from normal controls, while AstV-ND-3 [frontal cortex tissue]) clustered with frontal cortex from normal controls, validating this approach (Supplemental figure 5). AstV-ND-1 (brain tissue from unknown site) clustered with brainstem and medulla from normal controls. Additional comparison of the AstV-ND-1 sample of “unknown site” was made against all normal control samples representing different brain sites. The pairs of AstV-ND cases and normal control samples that were selected to identify DEGs for this study are indicated in Supplemental figure 5.

The metadata and normalized data files have been deposited in the GEO database (https://www.ncbi.nlm.nih.gov/geo/query/acc.cgi?acc=GSE201384).

**Supplemental Figure 5.**
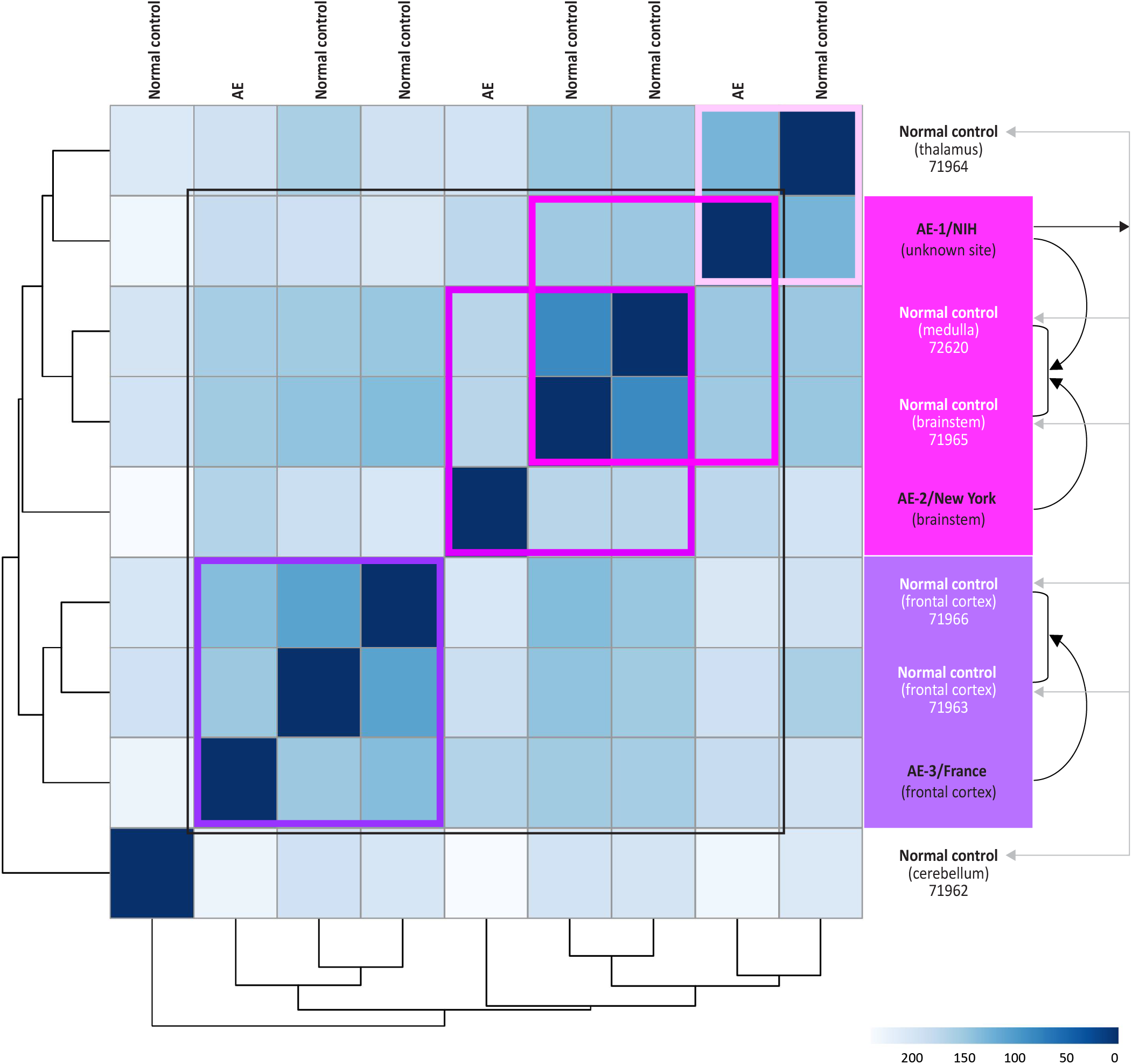
Clustering analysis of AE and normal control brain samples. The heatmap of the sample-to-sample distances in the transformed reads count matrix (dseq2). Samples that clustered together are highlighted by the same colors on the right and within the matrix. The pairs of AstV-ND and normal control samples chosen to identify DEGs for this study are shown on the right by arrows connecting relevant samples. Major comparisons for this study are shown by black curved arrows and outlined by a black square in the matrix. Additional comparison of the AstV-ND-1 sample of “unknown site” was made against all normal control samples representing different brain sites (gray arrows) (described later in the results). Close position of AstV-ND-1 unknown site to the normal control thalamus is outlined by a pink box.

### Genomic analyses

Functional enrichment analyses of gene expression data were performed using the PANTHER statistical enrichment test (SET) and gProfiler as previously described [21]. A multi-query multi-source gProfiler platform [18] was used for simultaneous comparative functional analyses of differential gene expression in three patients with AstV-ND cases (AstV-ND-1, AstV-ND-2, and AstV-ND-3) and in patients with HIV-1-associated neurocognitive disorders (HAND; 43-58 years old; frontal lobe white matter tissue; DEGs averaged from five postmortem samples; Table S2 in reference [17]). G:GOSt algorithm was used for multiple testing correction (https://biit.cs.ut.ee/gprofiler/page/docs).

### Astrovirus RNA detection

#### Virus microarray

RNA was isolated from frozen brain tissue sample from AstV-ND-1 case and hybridized to a microarray containing >3000 viral probes for viruses spanning 31 virus families known to infect vertebrates and >19,000 probes for endogenous genes. The virus microarray and bioinformatics analysis were performed as previously described [42]. An average probe intensity from control brain tissue was used to define the background threshold (average probe intensity plus three standard deviations).

#### PCR

RNA was isolated from frozen brain tissue sample from AstV-ND-1 case and cDNA was generated using a Superscript First-Strand Synthesis System for RT-PCR using random hexamers (Invitrogen). A first round of PCR was performed using primers panAV-F11 (5′-GARTTYGATTGGRCKCGKTAYGA-3′), panAV-F12 (5′-GARTTYGATTGGRCKAGGTAYGA-3′) and panAV-R1 (5′-GGYTTKACCCACATICCRAA-3′), followed by a second round of PCR with primers panAV-F21 (5′-CGKTAYGATGGKACKATICC-3′), panAV-F22 (5′-AGGTAYGATGGKACKATICC-3′) and panAV-R1 as previously described [43].

#### In situ hybridization

RNA was isolated from frozen brain tissue sample from AstV-ND-1 case and cDNA was generated using a Superscript First-Strand Synthesis System for RT-PCR using random hexamers (Invitrogen). Astrovirus capsid primers, Astrocap for 5- CGC GCG GAT CCA CCA TGG GGG GAT GGT GGT TTG TCA AG -3 and Astrocap rev 5- CGC GCG TCT AGA CTC GGC GTG GCC TCG GCG CAA -3 yielded a 1,227 bp band, which was cloned into pXLE42·V5 tag vector at the Bam HI and XbaI sites. After confirming the astrovirus capsid sequence, the negative strand was used instilling a 19-point bp change for probe creation, identical to the patient’s bp changes. Sections from the AstV-ND-1 patient’s brain and a control human brain were mounted onto microscope slides and hybridized with either astrovirus or actin probes using a QuantiGene (R superscript circularized) ViewRNA ISH Tissue 2-Plex Assay kit (Affymetrix Cat No. QVT0012).

The accession number for the astrovirus AstV-ND-1 sequence in Genbank is XXXXX.

### Astrovirus phylogenetic analysis

A search of the NCBI protein sequence database for Astroviridae ORF2 protein sequences identified 96 protein sequences. These sequences plus the translation of the HAstV-NIH sequence were aligned using the MUSCLE local alignment program [44] and this multiple sequence alignment was inspected and manually improved. Two different algorithms were used to calculate phylogenetic trees from this protein multiple sequence alignment. First, a neighbor-joining analysis [45] was performed using the algorithm implemented in the MEGA6 software [46] using the Jones-Taylor-Thorton substitution model [47] with rate variation among sites modeled by a discrete gamma distribution (shape parameter = 1.2). Support for individual clades was assessed using 500 bootstrap replicates [48]. Next, a Bayesian phylogenetic analysis was performed using the MrBayes software [49]. A single chain analysis running for 2,000,000 generations with sampling every 200 generations was performed. The program evaluated the best fit substitution model and chose the WAG model [50] with posterior probability = 1.0. All convergence criteria were met at the end of this analysis. No significant difference was found between the two phylogenies, so the Bayesian phylogeny was used going forward.

### Histology and Immunohistochemistry

FFPE brain tissue sections were stained with hematoxylin and eosin (H&E) or processed for immunohistochemistry. Brightfield colorimetric immunohistochemistry was performed using Bond RX (Leica Biosystems) according to manufacturer protocols. Diaminobenzidine was used for colorimetric detection (brown) and hematoxylin was used for counterstaining. The following primary antibodies were used for brightfield colorimetric immunohistochemistry: antibody against astrovirus capsid protein [6, 10] (rabbit polyclonal; 1:500), anti-CD68 (mouse monoclonal [KP1]; Biocare Medical; 1:50), anti-GFAP (rabbit polyclonal; Abcam; 1:250), anti-CD3 (rat monoclonal [Clone 12]; AbD Serotec; 1:600); anti-CD20 (mouse monoclonal [Clone L26]; Agilent; 1:200); anti-VGLUT1 (rabbit polyclonal; Synaptic Systems; 1:300), and anti-VGAT (rabbit polyclonal; Synaptic Systems; 1:300). Double immunofluorescent staining was performed using Bond RX (Leica Biosystems) according to manufacturer protocols with the following primary antibodies: antibody against astrovirus capsid protein [6, 10] (rabbit polyclonal; 1:500); anti-MAP2 (mouse monoclonal [Clone 198A5]; Synaptic Systems; 1:300), and anti-synaptophysin (mouse monoclonal [SY38]; Abcam; 1:10). Host appropriate secondary antibodies were labeled with a red fluorescent dye Alexa Flour 594 (Life Technologies; 1:300) or biotinylated secondary antibody (Vector Laboratories; 1:200) and green fluorochrome streptavidin 488 (Life Technologies; 1:500). Detection was completed using the Bond Research Detection kit (Leica Biosystems; CAT# DS9455). Nuclei were counterstained with DAPI (Vector Laboratories), and sections were mounted with ProLong Gold anti-fade reagent (Invitrogen). EverBrite TrueBlack mounting medium (Biotium) was used to attempt to quench the lipofuscin autofluorescence according to manufacturer protocol with limited success.

### Image acquisition and analysis

Brightfield colorimetric immunohistochemistry and H&E-stained sections were scanned at x40 magnification using the ScanScope AT2 (Leica Biosystems), and ImageScope software was used for digital slide analysis. Mantra Imaging System and InForm software (Akoya Biosciences) were used for multispectral immunofluorescent image acquisition, spectral unmixing, and generation of simulated H&E images based on tissue autofluorescence according to manufacturer protocols. Simulated H&E images based on tissue autofluorescence were used as a tissue topographical reference for immunofluorescent images and to aid in identification of autofluorescent lipofuscin granules in corresponding immunofluorescent images.

## Acknowledgements

This work was supported by the NIAID Intramural Research Program. We thank Melissa Pasquale-Styles in the Office of Chief Medical Examiner, City of New York, and Dr. Marc Eloit in the Institut Pasteur, Paris, France for assistance with tissue samples used in this research. We thank Dr. Stefano Morenco in the Human Brain Collection Core, Intramural Research Program, NIMH (http://www.nimh.nih.gov/hbcc) for providing brain tissue samples from subjects without known neurological disease that were used as normal controls in this research.

This work was supported by the NIAID Intramural Research Program.

